# Helminth infection induces neuroimmune remodeling and clinical remission in a mouse model of multiple sclerosis

**DOI:** 10.1101/2025.09.02.672458

**Authors:** Naomi M Fettig, Sarah J Popple, Madilyn B Portas, Ava J Sharon, Arman Sawhney, Thomas Worthington, Blair K Hardman, Morgan Coburn, Ukpong B Eyo, Mark C Siracusa, Marc S Horwitz, Lisa C Osborne

## Abstract

The central nervous system (CNS) is under constant immunosurveillance and influenced by immune-related effector molecules, including type 2-associated cytokines. Long-lasting type 2 immunity elicited by intestinal helminth infections can modify immune responses and wound repair locally and in peripheral tissues, but direct effects of helminth infection on the CNS are poorly understood. Here, we explore whether naturally-evoked type 2 immune responses can modify neuroimmune interactions for therapeutic gain in a mouse model of multiple sclerosis. Chronic infection with the helminth *Trichinella spiralis* (*Ts*) remodelled the neuroimmune landscape, including establishment of a robust population of CNS-resident T helper 2 cells, which subsequently minimized CNS inflammation and demyelination during experimental autoimmune encephalomyelitis (EAE). Clinical remission could be achieved with prophylactic or therapeutic infection, was *Stat6*-dependent, and adoptive transfer of Th2 cells promoted remission in the absence of overt infection. These findings highlight the potential for harnessing type 2 immunity to modify outcomes of neuroinflammation and neurodegeneration.

**Summary:** Fettig *et al.* demonstrate that infection with the helminth *Trichinella spiralis* elicits rapid recruitment and sustained presence of Th2 cells in the central nervous system where they modify microglia function and are implicated in resolving autoimmune-mediated paralysis and neuroinflammation.

## Introduction

Multiple sclerosis (MS) is a chronic autoimmune disease of the central nervous system (CNS) characterized by infiltration of autoreactive T cells, resulting in demyelination and degeneration of neurons. Highly efficacious disease-modifying therapies for relapsing-remitting MS (RRMS) generally target aspects of lymphocyte activation or trafficking into the CNS to prevent inflammation and myelin damage. However, these immunomodulatory therapies are largely ineffective in progressive stages of MS, and a knowledge gap in our understanding of fundamental mechanisms of disease progression in MS have hindered development of treatments that prevent, halt, or treat progression (1). In RRMS, the cyclical nature of clinical symptoms and of lesioning in the CNS can be at least in part explained by a shifting equilibrium of demyelinating and remyelinating processes (2). One hypothesis to explain the switch from RRMS to progressive MS is a cessation of remyelination, which results in a lack of trophic support for neurons and leaves neurons susceptible to degeneration and cytotoxic attack (3-5). Thus, therapeutic strategies to promote remyelination or repair tissue damage in the CNS represent the next frontier in disease-modifying therapy for progressive MS.

The CNS has historically been regarded as an immune-privileged niche that lacks substantial surveillance by leukocytes. However, emerging evidence demonstrating roles for CNS-infiltrating leukocytes in neurodevelopment (6, 7), homeostatic immune surveillance (8-12), and resolution of inflammation (13, 14), establishes an opportunity to define and harness pathways of tissue-protective immunity as a therapeutic strategy for neurological diseases. In peripheral tissues such as the skin and intestine, type 2 immune cells such as T helper 2 (Th2) cells, type 2 innate lymphoid cells (ILC2s), and alternatively activated macrophages have well-established pro-reparative functions (15, 16), and the contributions of these cells to resolution of CNS pathology are beginning to be appreciated. Indeed, there is a growing body of evidence for type 2 response-initiating alarmins and type 2 effector cytokines in neuroprotection or repair in models of CNS injury, stroke, and mouse models of MS (17). However, utilization of these molecules has yet to be translated into the clinic, highlighting an ongoing need to uncover neuro-modulatory mechanisms of type 2 immunity in neuroinflammatory disease.

Helminths are multicellular macro-parasites that co-evolved with the immune system. The classic type 2 immune responses activated in response to helminth infection balance pathogen clearance while prioritizing tissue homeostasis (18, 19). Without minimizing the burden of helminth infections across the globe, the “Old Friends” hypothesis suggests that helminth infections provide important immune-educating stimuli, and that lack of such exposures increases the risk of hypersensitivity to otherwise innocuous stimuli that can lead to allergy or autoimmunity (20, 21). In the context of ongoing inflammation, infection with helminths could re-balance the immune setpoint in a manner that resolves inflammation and favors tissue repair (22). To this end, helminth immunotherapy (HIT) has been explored as a method to dampen Th1- and Th17-driven inflammation and promote tissue repair in a variety of inflammatory and autoimmune diseases, including inflammatory bowel disease (23), allergy (24, 25), and rheumatoid arthritis (26, 27). While helminth infections have shown some prophylactic or therapeutic benefit in rodent models of MS (28), the few clinical trials of HIT in MS have reported limited success (29-32), and mechanistic insights from rodent HIT studies have yet to be translated for therapeutic benefit in people with MS.

The majority of studies investigating HIT in rodent models of MS focus on peripheral immunoregulation, leaving a substantial gap in our understanding of how helminths modulate immunity and glial cell function within the CNS during disease. To harness the reparative functions of helminth-elicited type 2 immunity in neuroinflammatory and neurodegenerative disease, pre-clinical studies of HIT must investigate perturbations to the site of inflammation and damage – the CNS – to identify pathways or immune mediators that may provide novel therapeutic strategies.

Elements confounding the translation of mechanistic understanding of helminths as immune modulators from rodent models to humans include the diversity in helminth species that have been trialed for HIT in humans and rodent models, the strict species-specific host tropism of many helminths, and variation in the immune-modulatory or immune-evasive strategies employed by these different helminths in their natural host environment (33-35). Studying immunomodulatory mechanisms of true “Old Friends” – that is helminths with natural human hosts – may provide the most translationally-relevant understanding of mechanisms of helminth-mediated immune modulation due to coevolution between helminths and host immune systems (36). *Trichinella spiralis* (*Ts*) is one such helminth with both human and mouse tropism that has coevolved over millennia to be one of the most widespread helminth infections (37-42).

Here, we identify an unappreciated neuroimmune-modulatory pathway elicited by infection with *Ts,* which promotes rapid infiltration and chronic persistence of neuroprotective leukocytes in the CNS to promote recovery from experimental autoimmune encephalomyelitis (EAE), a mouse model of MS. During acute *Ts* infection, type 2-polarized and regulatory immune cells infiltrate the brain and spinal cord and activate local microglia. Following transient *Ts-*mediated blood-brain barrier breakdown, *Ts-*reactive Th2s migrate to the CNS to establish a long-lived tissue-resident memory population, which later drives recovery from EAE. *Ts-*infected mice have a profound delay in the onset of EAE, which unlike the recovery phenotype, is independent of *Ts-*primed STAT6-dependent Th2s in the periphery. Importantly, *Ts* infection can be used therapeutically to prevent clinical relapses and limit CNS inflammation in a relapsing-remitting EAE model. Altogether, these findings deepen our understanding of the neuro-modulatory functions of *Ts* both at steady-state and during neuroinflammatory challenge and highlight mechanisms by which *Ts-* mediated immune modulation could be harnessed for HIT.

## Results

### Chronic *Ts* infection delays EAE onset and promotes recovery

To define the impact of persistent *Ts* infection on EAE, C57BL/6J female mice were infected with 400 *Ts* larvae (L1) by oral gavage and immunized with MOG_35-55_/CFA 4 weeks later (28 days post-infection) (**Figure 1A**). Consistent with results from other helminth infection models (43-45), disease onset was significantly delayed (mean ± SD day to onset: not-infected EAE, day 10.4 ± 2.6; *Ts/*EAE day 21.1 ± 5.0), and overall disease burden was diminished in *Ts-*infected mice compared to not-infected (NI) EAE controls (**Figure 1B-D, S1A**). Stratifying clinical disease outcomes of *Ts/*EAE mice by the extent of delayed disease onset unmasked features of altered clinical EAE trajectory in *Ts-*infected mice: although peak severity was similar between NI/EAE and *Ts/*EAE animals, *Ts-*infected mice reliably recovered motor function while paralysis persisted in NI/EAE animals (**Figure 1D, S1B**). To account for the variability in the time to disease onset, recovery metrics were calculated based on clinical EAE scores at the peak of disease and the 10-day post-peak interval for individual mice. This analysis demonstrated a more pronounced magnitude and rate of recovery in *Ts-*infected mice than uninfected EAE mice (**Figure 1E-G**), leading to an overall reduction in cumulative disease burden in the 10-day post-peak window (**Figure 1H**). We defined EAE “remission” as normalization of gait and limb function and recovery of tail tonicity (clinical score ≤ 5 on a 16-point scale) and determined that approximately 65% of *Ts-*infected mice met these criteria within 10 days post-peak EAE severity, as opposed to only 15% of uninfected mice (**Figure 1I**). Raw clinical data of all mice included in remission quantifications is included in **Table S1**. In alignment with the recovery of EAE symptoms, the spinal cord (SC) of *Ts/*EAE mice had reduced cellular infiltration and lesion formation (**Figure 1J-K, S1C)**, and more intact myelin compared to uninfected EAE controls 10 days after peak EAE (**Figure 1L-M, S1D**).

**Figure 1.**
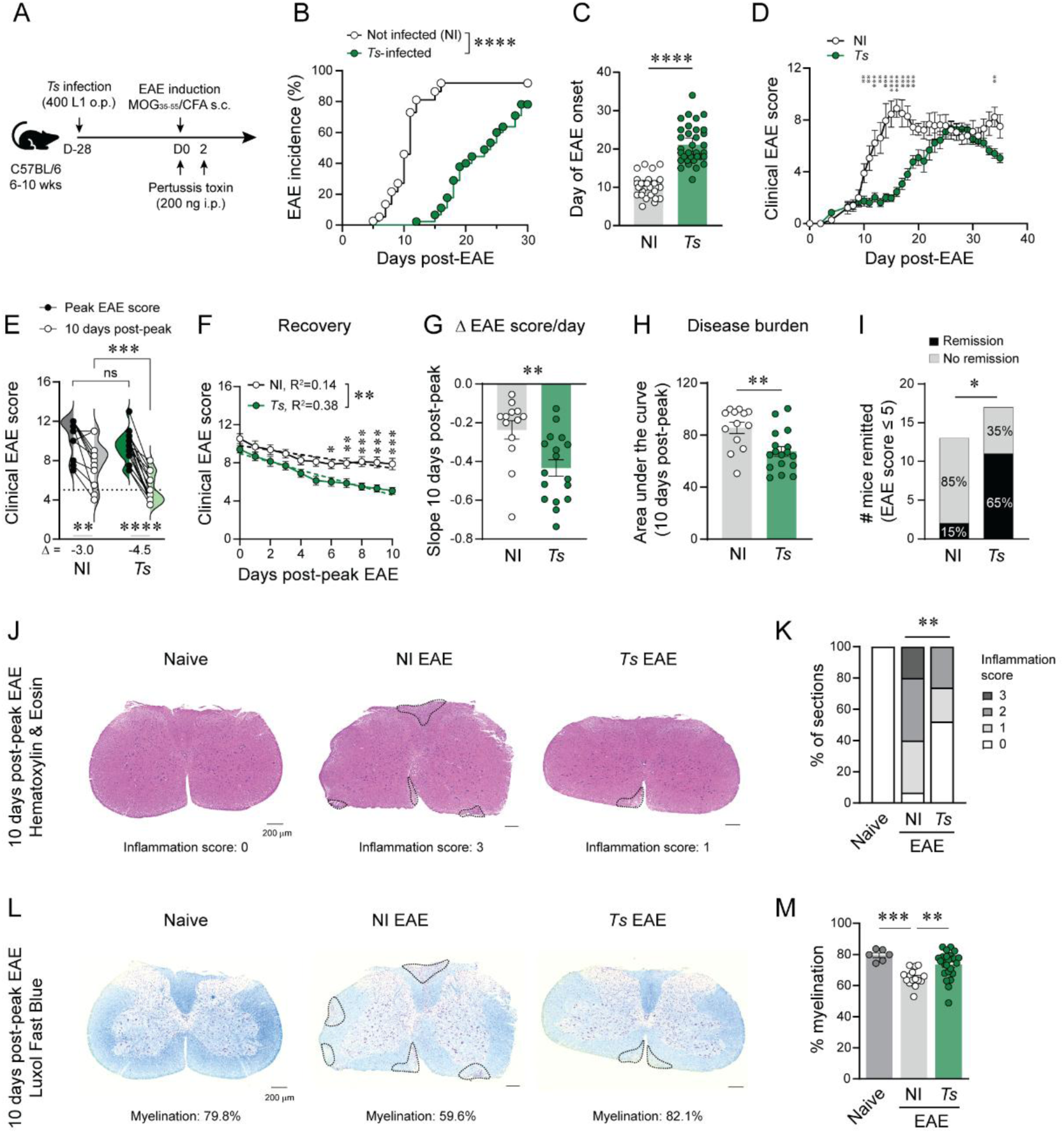
Chronic *Ts* infection delays EAE onset and promotes recovery. (A) Experimental design for *Ts/*EAE experiments. C57BL/6 mice were infected by oral gavage with 400 *Ts* L1. 4 weeks post-infection, EAE was induced by s.c injection of MOG_35-55_ in complete Fruend’s adjuvant (CFA) and i.p. injections on d0 and d2 of pertussis toxin. (B) EAE incidence of *Ts-* infected (n=45) or non-infected (NI: n=37) mice. (C) Quantification of the day of EAE onset. (D) Clinical EAE scores of NI (n=9) or chronically *Ts-*infected mice (n=18). Data pooled from 4 independent experiments, including only mice with clinical data to day 35 post-EAE. (E) Paired analysis of the peak EAE score reached vs EAE score 10 days after peak disease (NI: n=13, *Ts*: n=17). Overlaid violin plots depict data distribution. Delta (Δ) = median change in EAE score. (F) EAE scores 10 days post-peak disease. Dashed lines depict linear regression. (G) Slope of the linear regression for individual mice 10 days post-peak EAE. (H) Area under the curve for the 10-day post-peak interval. (I) Number of mice remitted (EAE score ≤ 5). (J) Representative H&E-stained spinal cord (SC) sections 10 days post-peak EAE. (K) Proportion of SC sections with a given “inflammation score” (scored from 0-4), compiled from 3 sections per SC (naïve (n=2), NI EAE (n=5), *Ts* EAE (n=8)). (L) Serial LFB-stained SC sections. (M) Percent myelinated area. Statistics by Log-rank (Mantel-Cox) test (B), Mann-Whitney test (C, G, H), multiple Mann-Whitney tests with Benjamini, Krieger, and Yekutieli multiple comparisons test (FDR=1%) (D), 2-way ANOVA with Tukey’s multiple comparisons test (E), 2-way ANOVA with Sidak’s multiple comparisons test and ANCOVA for linear regression slopes (F), Fisher’s exact test (I, K), or one-way ANOVA with Tukey’s multiple comparisons test (M).

### *Ts*-infected EAE mice have impaired Th17 priming

To interrogate whether delayed EAE onset in *Ts-*infected mice was due to altered CD4^+^ T cell activation, we evaluated the effect of *Ts* infection on pro-inflammatory and myelin-specific CD4^+^ T cells at day 9 post-immunization, a timepoint associated with systemic T cell activation in the secondary lymphoid organs prior to robust CNS infiltration and EAE onset. Independent of EAE, the frequency of antigen-experienced (CD44^hi^) CD4^+^ T cells was elevated in the spleen and inguinal lymph node (iLN) of *Ts-*infected mice (**Figure 2A**). However, EAE-induced expansion of the MOG_35-55_/CFA-draining iLN was impaired in *Ts/*EAE compared to NI/EAE, including fewer activated CD44^hi^ CD4^+^ T cells (**Figure S2A, B**). Further, the proportion of activated CD4^+^ T cells expressing the Th17-defining transcription factor RORγt was diminished in the spleen and iLN of *Ts/*EAE mice (**Figure 2B, C**), which was associated with a reduced fraction of IL-17A-expressing splenic CD4^+^ T cells and a trending decrease in the amount of cytokine expressed per cell (∼30% reduction in median fluorescence intensity (MFI), **Figure 2D-F**). Expression patterns of the Th1 differentiation markers TBET and IFNγ were not consistent across iLN and spleen (**Figure 2E, S2C**), a result that could be due to polyclonal analysis that does not distinguish between CD4^+^ T cells primed by *Ts* or by MOG_35-55_ immunization. To address this, splenocytes were stimulated with MOG_35-55_ peptide and the proportions of IFNγ-, GM-CSF-, and IL-17A-expressing CD4^+^ T cells quantified. Notably, pre-existing *Ts* infection led to reduced proportions of GM-CSF- and IL-17A-, but not IFNγ-producing MOG-specific CD4^+^ T cells (**Figure 2G, H, S2D**), along with a decrease in the MFI of IL-17A (**Figure 2I**). Consistent with these data suggesting an impairment in Th17 polarization of MOG-reactive cells, there was a reduction in the proportion and number of RORγt^+^ MOG I-A^b+^ CD4^+^ T cells in both the spleen and iLN (**Figure 2J, K)**, although the total number of MOG I-A^b+^ CD4^+^ T cells was reduced only in the iLN (**Figure S2E**).

**Figure 2.**
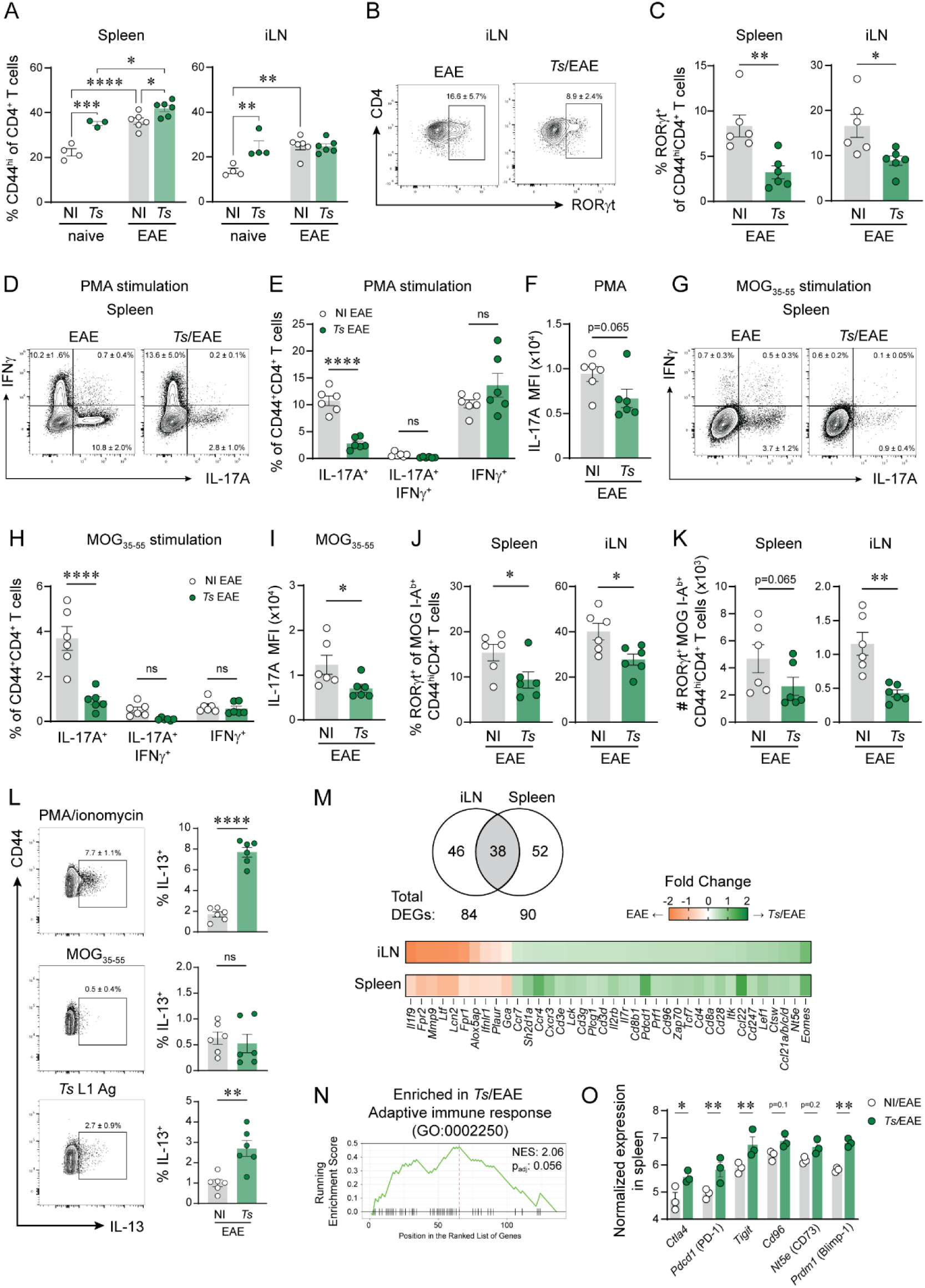
*Ts* alters immune priming in EAE. Spleens and inguinal lymph nodes (iLNs) were harvested on day 9 post-EAE (or non-EAE naïve controls) from chronically *Ts-*infected or non-infected (NI) mice. (A) Proportion of CD44^hi^ CD4^+^ T cells in the spleen or iLN. (B) Concatenated flow plots (n=6 per group) of RORγt expression in CD44^hi^CD4^+^ T cells in the iLN. (C) Proportion of RORγt^+^ CD44^hi^CD4^+^ T cells in the spleen and iLN. (D) Concatenated flow plots (n=6 per group) of splenocytes stimulated with PMA/ionomycin, gated on CD44^hi^CD4^+^ T cells. (E) Proportion of IL-17A^+^, IFNγ^+^, or IL-17A^+^IFNγ^+^ CD44^hi^CD4^+^ T cells, and (F) median fluorescence intensity (MFI) of IL-17A by IL-17A^+^CD44^+^ CD4^+^ T cells following PMA/ionomycin stimulation. (G) Concatenated flow plots (n=6 per group) of splenocytes stimulated with MOG_35-55_, gated on CD44^hi^CD4^+^ T cells. (H) Proportion of IL-17A^+^, IFNγ^+^, or IL-17A^+^IFNγ^+^ CD44^hi^CD4^+^ T cells, and (I) MFI of IL-17A by IL-17A^+^CD44^+^ CD4^+^ T cells following MOG_35-55_ stimulation. (J) Proportion and (K) number of RORγt^+^ MOG_38-49_ I-A^b+^ CD44^hi^CD4^+^ T cells in the spleen and iLN. (L) Concatenated flow plots (left) and quantification (right) of IL-13 expression in CD44^hi^CD4^+^ T cells (n=6 per group) from splenocytes stimulated with PMA/ionomycin (top), MOG_35-55_ (middle), or *Ts* L1 antigen (Ag, bottom). (M) Nanostring nCounter of bulk iLN and spleen RNA. Venn diagram shows the number of differentially expressed genes (DEGs) in the iLN, spleen, or both in *Ts/*EAE vs NI/EAE. Heatmap shows 35 common DEGs in spleen and iLN. (N) GSEA of combined spleen and iLN gene lists. (O) Normalized expression of select genes in the spleen. Statistics by 2-way ANOVA with Tukey’s multiple comparisons test (A), unpaired t-tests (C, F, I, J, L), 2-way ANOVA with Sidak’s multiple comparisons test (E, H, O), or Mann-Whitney test (K). Data inset in flow plots show mean ± SD.

Regulatory CD4^+^ T cells (Tregs) can inhibit immune priming following MOG_35-55_ immunization (46-48). Independent of EAE, *Ts*-infected mice had an increased proportion of Tregs (FOXP3^+^ CD4^+^) in the secondary lymphoid organs compared to uninfected naïve controls (**Figure S2F**). Following EAE induction, however, this *Ts-*mediated enhancement was lost in the spleen and significantly reduced in the iLN. Further, the number of MOG I-A^b+^ Tregs cells was similar between NI/EAE and *Ts/*EAE mice (**Figure S2H**), suggesting that regulatory myelin-reactive cells were not generated in place of classical inflammatory CD4^+^ effectors in *Ts*/EAE. Even with a similar proportion of Tregs in the spleen, the ratio of effector T cells (IFNγ and/or IL-17A-producing) to FOXP3^+^ Tregs was significantly reduced in *Ts/*EAE, following either PMA or MOG_35-55_ stimulation (**Figure S2G**), which may suggest a role for Tregs in limiting expansion of effector T cells in response to MOG_35-55_ immunization.

*Ts/*EAE splenocytes produced the Th2-associated cytokine IL-13 following polyclonal stimulation, but there was negligible expression of IL-13 following MOG_35-55_ peptide stimulation in either NI/EAE or *Ts/*EAE splenocytes. Stimulation with *Ts* L1-derived antigen, however, did promote IL-13 production by *Ts/*EAE CD4^+^ T cells (**Figure 2L**). These data indicate that a MOG-reactive Th2 population is not generated in *Ts/*EAE mice. Instead, a residual *Ts*-reactive Th2 population remains in the spleen, highlighting the persistent nature of *Ts-*reactive cells in secondary lymphoid organs (SLOs).

We evaluated the transcriptional profile of NI/EAE and *Ts/*EAE spleens and iLNs using the Nanostring nCounter Mouse Host Response panel, which contains probes for over 700 immune-associated genes. We identified 62 upregulated and 28 downregulated genes (fold change >1.2, p<0.05) in *Ts/*EAE spleens compared to NI/EAE spleens, and 51 upregulated and 33 downregulated genes in *Ts/*EAE iLNs compared to NI/EAE iLNs (**Table S2**, **Figure S2I**). Of the differentially expressed genes (DEGs), 38 were shared in the spleen and iLN, always with the same directionality between tissues (**Figure 2M**). In alignment with the observed increase in T cell activation (CD44^hi^) by flow cytometry in *Ts/*EAE SLOs, T cell activation genes were enriched in both SLOs of *Ts/*EAE mice, including increased expression of genes involved in TCR signaling (*Cd3e/d/g, Cd247, Cd4, Cd8a/b, Cd28, Zap70, Lck, Itk, Plcg1*) and the formation of memory T cells (*Tcf7, Il7r, Eomes, Lef1, Ccr7*). Collectively, *Ts/*EAE SLOs were enriched for genes associated with the adaptive immune response (GO:0002250) by gene set enrichment analysis (GSEA) (**Figure 2N**). Along with increased T cell activation, the *Ts/*EAE spleen had an increased transcriptional signature associated with inhibitory co-receptor and immune checkpoint expression, including *Pdcd1* (PD-1), *Ctla4, Tigit, Cd96, Nt5e* (CD73) and *Prdm1* (Blimp-1) (**Figure 2O**). Only *Pdcd1* (PD-1), *Nt5e* (CD73), and *Cd96* were significantly increased in the *Ts/*EAE iLN (**Figure S2J**).

Upregulated genes in NI/EAE mice trended towards an innate immune signature, including increased expression of *Myd88, Irak3, Il1b, Il18, Ccr2, Cxcl5, C5ar1, Csf3r, Fcgr2b, Fcgr4,* and *Ms4a4a* in the iLN; increased expression of *Itgam, Lta4h, Elane, Ncf2,* and *Peli2* in the spleen; and *Mmp9, Alox5ap, Fpr1/2, Ifnlr1, Gca, Ltf,* and *Lcn2* in both SLOs. A few notable exceptions were upregulated in *Ts/*EAE spleens, including *Irf4,* a negative regulator of cytokine production following TLR stimulation (49), and *Xbp1*, which is associated with promotion of the Th2 response and an immunosuppressive state in DCs (50, 51). These data suggest that myeloid cell activation and function is altered in the SLOs of *Ts/*EAE mice, which may impact the ability of T cells in these organs to receive optimal Th1/17 activation and polarization signals for development of EAE. This hypothesis aligns with previously published work that identified a role for *Ts* in inhibition of DC-mediated T cell priming (52-55).

Together, these data highlight a defect in the ability of CD4^+^ T cells to be activated towards a pro-inflammatory Th17 phenotype following immunization with MOG_35-55_/CFA, despite having a highly activated basal state. The impaired generation of myelin-reactive Th17s may impact the ability of these cells to initiate neuroinflammation.

### *Ts* alters effector T cell and microglia responses during peak EAE

Next, we investigated how alterations to the myelin-reactive CD4^+^ T cell pool during disease initiation could impact the neuro-inflammatory environment. During chronic EAE (day 40 post-immunization), the CNS of *Ts/*EAE mice had fewer pro-inflammatory RORγt^+^ Th17s but increased proportions and numbers of type 2 immune cells in the CNS, including GATA3^+^ Th2s and ILC2s (**Figure S3A, B**). These trends were consistent in the dural meninges (**Figure S3C, D**). However, cellular infiltration and cytokine production by CNS-infiltrating cells are greatest during acute EAE, which recede as disease becomes chronic (56, 57). This phenomenon was reproduced in our hands, whereby the time since EAE onset was negatively correlated with both the number of CNS-infiltrating leukocytes (CD45^hi^) and the proportion of pro-inflammatory cytokine-producing CD4^+^ T cells (IFNγ^+^ and/or IL-17A^+^) in the brain and spinal cord of NI/EAE mice (**Figure S3E, F).**

To account for differences in EAE onset, we evaluated effector T cell functions in the CNS of *Ts/*EAE and NI/EAE mice near the peak of their EAE disease. To this end, CNS tissue was collected at the peak of disease (day 13 post-EAE for uninfected mice, and days 23 or 36 post-EAE for *Ts-*infected mice), when all mice had a similar clinical EAE severity for similar lengths of time (**Figure S3G, H**). At peak EAE, total leukocyte infiltration into the CNS was not significantly different between NI/EAE and *Ts/*EAE (**Figure S3I**); however, the number of CNS-infiltrating CD4^+^ T cells, the canonical disease-initiating cells in EAE, was significantly reduced in *Ts/*EAE mice (**Figure S3J**). The activation state of CNS-infiltrated CD4^+^ T cells was strikingly different: expression of IL-17A^+^ and/or IFNγ^+^ by CD4^+^ T cells was reduced in the *Ts/*EAE spinal cord following either polyclonal or MOG_35-55_-specific stimulation (**Figure 3A-C**). Unlike NI/EAE, CNS-infiltrating Th1/17 effector CD4^+^ T cells were not correlated with the time since EAE onset and instead remained low throughout the duration of EAE in *Ts/*EAE mice (**Figure S3F**). Total numbers of GM-CSF^+^ CD4^+^ T cells were similarly reduced in *Ts/*EAE compared to NI/EAE, although proportional expression of GM-CSF was similar (**Figure S3K**). Further, numbers and proportions of IFNγ-expressing CD8^+^ T cells were reduced in *Ts/*EAE mice, despite similar total numbers of CNS-infiltrated CD8^+^ T cells (**Figure S3J, L**).

**Figure 3.**
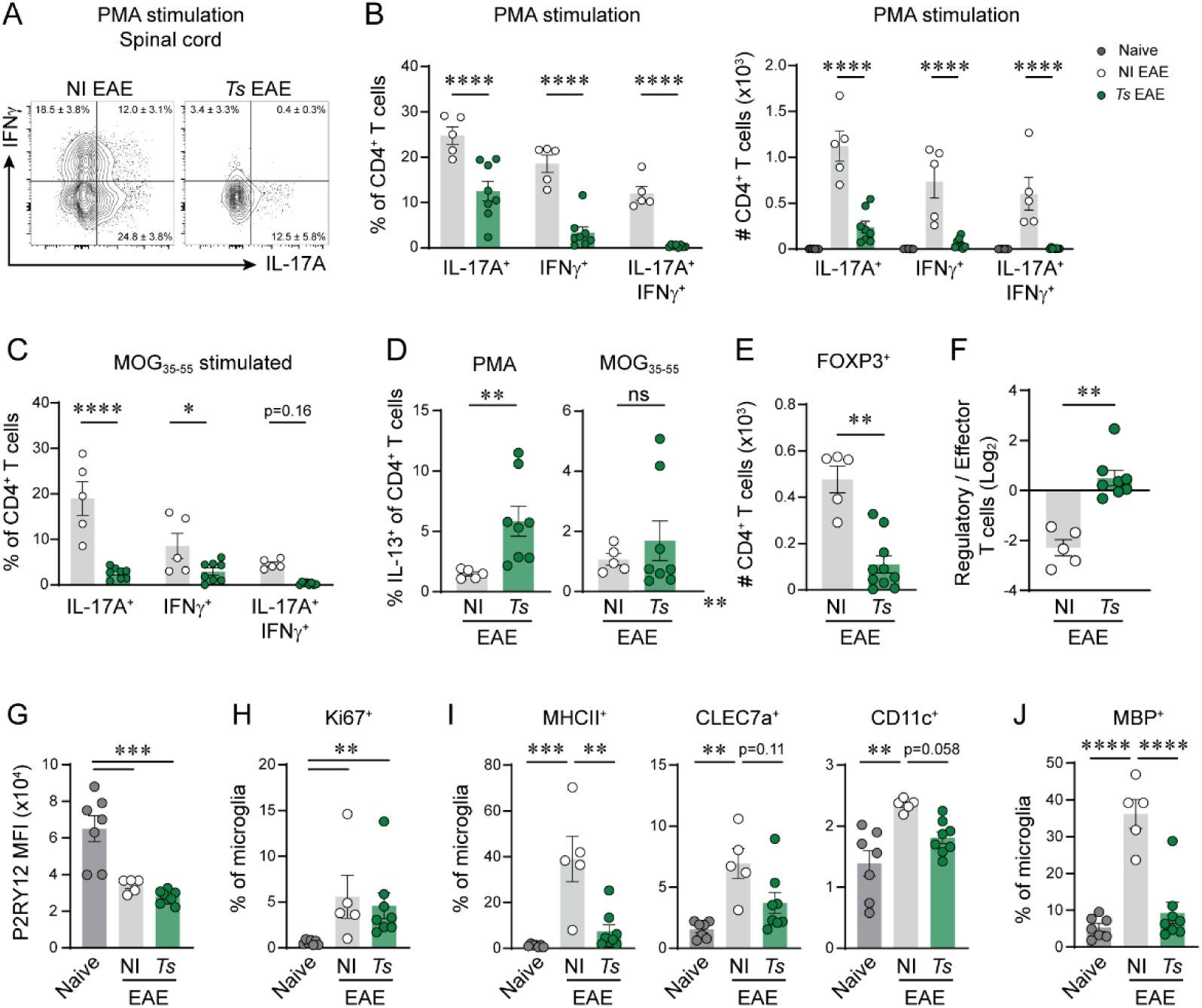
Chronic *Ts* infection alters effector T cell and microglia responses during peak EAE. CNS tissue was collected at peak EAE from *Ts-*infected or uninfected mice. (A) Concatenated flow plots of (n=5-8 per group) of IL-17A and IFNγ expression in CD4^+^ T cells from spinal cords (SC) stimulated with PMA/ionomycin. (B) Proportion (left) and number (right) of IL-17A^+^, IFNγ^+^, or IL-17A^+^IFNγ^+^ CD4^+^ T cells from PMA/ionomycin-stimulated SC. (C) Proportion of CD4^+^ T cells expressing IL-17A^+^, IFNγ^+^, or IL-17A^+^IFNγ^+^ following overnight MOG_35-55_ stimulation of SC. (D) Proportion CD4^+^ T cells that express IL-13 following PMA stimulation (left) or MOG_35-55_ stimulation (right) of SC. (E) Number of FOXP3^+^ CD4^+^ T cells from SC. (F) Log_2_-transformed ratio of the number of FOXP3^+^ Tregs to effector T cells (IL-17A^+^ and/or IFNγ^+^) in PMA-stimulated SC. (G) Median fluorescent intensity (MFI) of P2RY12 in microglia (CD45^mid^CD11b^+^P2RY12^+^) in brain. (H) Proportion of Ki67^+^ microglia. (I) Proportion of MHCII^+^, CLEC7a^+^, or CD11c^+^ microglia. (J) Proportion of microglia that contain intracellular MBP. Statistics by 2-way ANOVA with Šídák’s multiple comparisons test (B, C), Mann-Whitney test (D-F), one-way ANOVA with Tukey’s multiple comparisons test (G, I, J), or Kruskall-Wallis with Dunn’s multiple comparisons test (H). Data inset in flow plots show mean ± SD.

The proportion of IL-13-producing Th2s was increased in the spinal cord of *Ts/*EAE mice following polyclonal stimulation; however, similar to the priming phase of disease, IL-13^+^ CD4^+^ T cells in *Ts/*EAE spinal cords were not MOG_35-55_-reactive (**Figure 3D**). These data support the hypothesis that MOG-reactive CD4^+^ T cells are not polarizing towards a Th2 phenotype in place of the classical Th1/17 profile in EAE. MOG-reactive CD4^+^ T cells also did not appear to adopt a regulatory phenotype either, as the proportion of Tregs recovered following overnight MOG re-stimulation was similar in NI/EAE and *Ts/*EAE (**Figure S3M**), and the total number of Tregs was reduced in *Ts/*EAE spinal cords (**Figure 3E**). However, *Ts/*EAE spinal cords had a significantly increased ratio of FOXP3^+^ Tregs to effector T cells (IFNγ and/or IL-17A-producing) in the spinal cords following both polyclonal and MOG_35-55_ restimulation (**Figure 3F, S3N**); thus, Tregs could be dampening inhibiting inflammatory T cells *in situ* in the CNS during symptomatic EAE. Alternatively, since Tregs are typically generated in the recovery phase of EAE in response to inflammation in the CNS (58-60), the reduced number of Tregs may be a consequence of the minimal CD4^+^ T cell-mediated inflammation and demyelination in *Ts/*EAE, thereby reducing the need to induce Tregs in the CNS.

While EAE is considered a CD4^+^ T cell-mediated disease, myeloid cells are critical for disease initiation and damage of the myelin sheath. Microglia have complex multifaceted roles during EAE, including contributions to inflammatory damage and demyelination, as well as facilitation of myelin debris clearance which is crucial to resolving inflammation (61). Microglia from both NI/EAE and *Ts/*EAE mice were more activated and proliferative than naïve microglia by downregulation of P2RY12 and increased Ki67 expression (**Figure 3G, H**); however, microglia from *Ts/*EAE mice had significantly reduced expression of several antigen presentation and phagocytosis-associated markers such as MHCII, CLEC7a, and CD11c (**Figure 3I**). Further, fewer microglia contained intracellular myelin basic protein (MBP), indicating reduced phagocytosis of myelin (**Figure 3J**). In combination with reduced numbers of pathogenic effector T cells in the CNS and reduced demyelination, these data suggest that the myelin sheath is spared from demyelinating damage, resulting in reduced need for myelin debris clearance by microglia.

Together, these data suggest that MOG-primed CNS-infiltrating T cells in *Ts-*infected mice may be insufficient in function and/or number to cause severe demyelination and maintain severe disease. This reduced pro-inflammatory CD4^+^ T cell function could be due to impaired priming in the periphery or active inhibition in their effector function by the increased presence of type 2 cytokine-expressing Th2s or Tregs in the CNS.

### Acute *Ts* infection triggers remodeling of the neuroimmune landscape

Acute *Ts* infection has previously been shown to induce infiltration of neuroprotective monocytes into the brain (62), but the longevity of infiltration, or whether lymphocytes follow a similar migration pattern, remained unknown. Given the presence of non-myelin reactive Th2s in the CNS during EAE, we hypothesized that T cells were infiltrating the CNS following initial *Ts* infection, rather than during subsequent neuroinflammatory challenge. CD45^hi^ leukocytes infiltrated the CNS during acute *Ts* infection (10 DPI) and were maintained chronically (28 DPI) (**Figure S4A**). The number of myeloid cells (CD11b^+^) and T cells (TCRβ^+^) were increased at 10 DPI, but only T cells remained numerically and proportionally increased during chronic infection (**Figure 4A, S4B**). Consistent with previous findings (62), the acute increase in myeloid cells was driven by a variety of innate immune cells, including monocytes, macrophages, and DCs, among others (**Figure S4C**).

**Figure 4.**
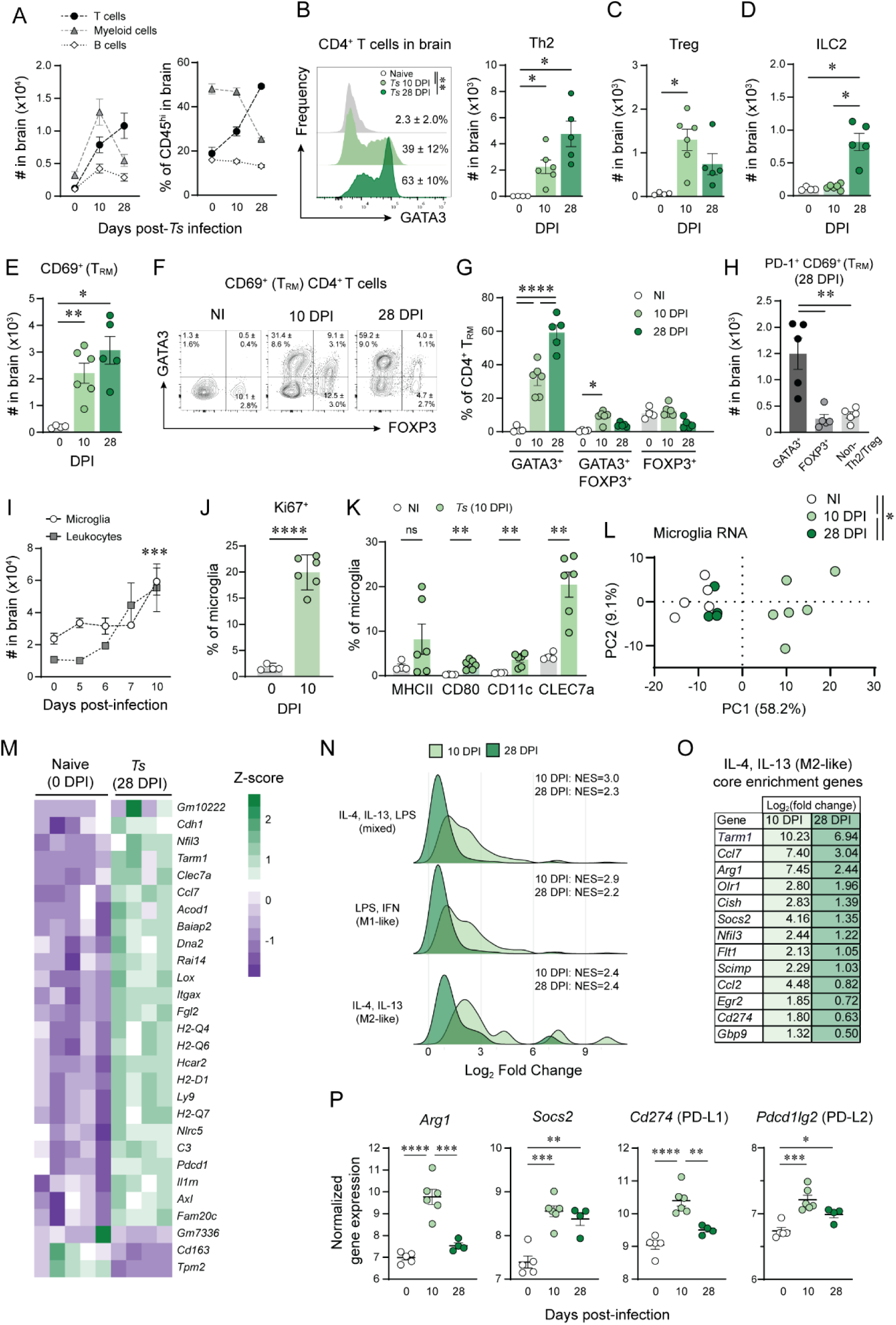
Acute *Ts* infection drives remodeling of the neuroimmune landscape. Mice were infected with *Ts* or left not infected (NI) and tissues collected at 10 DPI or 28 DPI. (A) Number of CD45^hi^ leukocytes in the brain and spinal cord. (B) Concatenated histograms (n=4-6) of GATA3 expression in CD4^+^ T cells in the brain (left) and quantification (right). (C) Number of FOXP3^+^CD4^+^ Tregs in the brain. (D) Number of ILC2s (GATA3^+^ TCRβ^-^B220^-^CD11b^-^CD11c^-^ CD45^hi^) in the brain. (E) Number of CD69^+^ CD4^+^ T cells (tissue resident memory, T_RM_) in the brain. (F) Concatenated flow plots (n=4-6) of GATA3 and FOXP3 expression, gated on CD69^+^CD4^+^ in the brain. (G) Percent of CD4^+^ T_RM_ in the brain that express GATA3 and/or FOXP3. (H) Number of PD-1^+^ CD4^+^ T_RM_ in the brain that express GATA3, FOXP3, or neither (non-Th2/Treg). (I) Number of microglia (CD45^mid^CD11b^+^P2RY12^+^) and leukocytes (CD45^hi^) in the brain during acute *Ts* infection. (J) Proportion of Ki67^+^ microglia. (K) Proportion of MHCII^+^, CD80^+^, CD11c^+^, or CLEC7a^+^ microglia. (L) Principal component analysis of microglia transcriptomes by RNA-sequencing. (M) Hierarchical clustering of all differentially expressed genes in microglia from 28 DPI vs naive mice. (N) Gene set enrichment analysis (GSEA) against published gene lists of M1, M2, or mixed culture conditions of bone marrow derived macrophages. (O) Core enrichment genes associated with enrichment of the M2-like dataset. (P) Normalized gene expression of select genes of interest. Statistics by one-way ANOVA with Tukey’s multiple comparisons test (A-E, H, P), two-way ANOVA with Tukey’s (G) or Dunnett’s multiple comparisons test (I), unpaired t-test (J, K), or PERMANOVA with multiple pairwise comparisons (L). ROUT outlier test (Q=1%) performed on data in (A) and data from one mouse removed from analysis. Data inset in flow plots show mean ± SD.

Within the T cell compartment, the proportion and number of CNS-infiltrated Th2s increased over the course of infection, making up a majority of the CD4^+^ T cells by 28 DPI (**Figure 4B, Figure S4D**). In contrast, the proportion and number of Tregs increased during acute infection but receded by 28 DPI (**Figure 4C, S4E**). ILC2s also increased in number, but only at the chronic timepoint evaluated (**Figure 4D**). In addition to cellular infiltration into the CNS parenchyma, the number and frequency of Th2s and Tregs, but not ILC2s, were increased in the dural meninges at 10 DPI (**Figure S4F-H**). The persistent nature of these CD4^+^ T cells could be explained by their high expression of the tissue residency marker CD69, most of which were GATA3^+^ Th2s (**Figure 4E-G**). Th2 and Treg tissue resident memory (T_RM_) cells were enriched for PD-1 expression (**Figure S4I**), resulting in a significant increase in the number of PD-1^+^ Th2 T_RM_ in the brain during chronic *Ts* infection (**Figure 4H**). PD-1 expression has been implicated in maintenance of tissue-resident Th2 populations (63), and PD-1:PD-L1 interactions have been shown to stabilize Th2 effector function (64), thus expression of chronic expression of PD-1 may contribute to the maintenance of Th2 T_RM_ in the CNS of *Ts-*infected mice.

To further investigate *Ts-*mediated alterations to the CNS neuroimmune landscape, we investigated the activation state of the local microglia population. Shortly after the appearance of leukocytes in the CNS, microglia began to proliferate (**Figure 4I-J**) and increased expression of select activation markers including MHCII, CD80, CLEC7a, and CD11c (**Figure 4K**). To develop an unbiased understanding of the microglia activation profile during acute and chronic infection, we performed RNA sequencing on FACS-purified microglia from naïve, acutely infected (10 DPI), and chronically infected (28 DPI) mice (**Figure S4J**). The overall transcriptional profile was significantly different between microglia from acutely infected and naïve mice (p=0.002, F=3.9), acutely and chronically infected mice (p=0.007, F=3.9), and chronically infected and naïve mice (p=0.02, F=1.4) (**Figure 4L**). Microglia from acutely infected mice were highly activated, with 935 differentially expressed genes (log_2_FC >1 or <1, p<0.05), including 608 upregulated and 327 downregulated genes; while microglia from chronically infected mice had only 29 differentially expressed genes, including 26 upregulated and 3 downregulated genes (**Table S3**).The relatively small number of differentially expressed genes at 28 DPI suggest a return to a near-baseline transcriptional state, but with some maintained alterations.

Several of the genes and proliferation markers observed by flow cytometry were similarly significantly upregulated at the transcriptional level during acute infection, including *Clec7a* (log_2_FC=2.8), *Itgax* (CD11c) (log_2_FC=2.5), *Mki67* (log_2_FC=3.7), and several MHCII gene loci such as *H2-Ab1* (log_2_FC=2.5). At 10 DPI, microglia downregulated several homeostatic genes (*P2ry12, Tmem119,* and *Sall1*), and the DEGs were significantly enriched for published gene lists associated with microglia priming, phagocytosis, and injury response by GSEA (**Figure S4K, L**) (65-67), altogether suggesting broad activation of microglia.

Of the genes that remained differentially expressed at 28 DPI, many indicate prolonged immunomodulation of microglia, including maintained upregulation of *Itgax* (CD11c), *Clec7a, Pdcd1* (PD-1), and of several non-classical MHCIb genes that have been associated with immune suppressive function and EAE resistance (68) (**Figure 4M**). Further, the microglial gene signature at 28 DPI maintained enrichment for activation- and phagocytosis-associated gene sets (**Figure 4L**). Other notable genes that remain upregulated in microglia at 28 DPI include *Axl, Hcar2, Nfil3,* and *Acod1,* which have neuroprotective roles in various models of neuroinflammation or neurodegeneration (69-74).

Using a published RNA-sequencing dataset of bone marrow-derived macrophages (BMDMs) cultured in M1-skewing conditions (LPS and IFNγ), M2-skewing conditions (IL-4 and IL-13), or mixed culture conditions (IL-4, IL-13, and LPS) (75), we evaluated if microglia from *Ts-*infected were enriched for genes associated with these classical polarization pathways. The 10 DPI microglia were highly enriched for all 3 gene sets, with the highest enrichment for the mixed culture conditions. All three gene sets remained significantly enriched in the 28 DPI microglia, but the M2-skewed gene list was the most enriched at the chronic timepoint and was the only condition that maintained a comparable enrichment score between the two timepoints post-*Ts* infection (**Figure 4N**). The core enrichment genes for the M2-skewed gene list included increased expression of *Arg1, Socs3,* and *Cd274* (PD-L1), among others (**Figure 4O-P**). *Pcdc1lg2* (PD-L2) similarly remained slightly upregulated at 28 DPI **(Figure 4P**), which can be induced by type 2 cytokine signaling similarly to macrophages (76, 77). In addition to PD-L1 interactions stabilizing Th2 responses (63, 78), PD-L1 may simultaneously inhibit activation of CNS-infiltrated Th1/17s or CD8 T cells during EAE (79-81). Sustained *Pdcd1* (PD-1) expression may also limit their activation through interaction with other PD-L1-expressing glial cells (82). These data suggest that while microglia from chronically infected mice return to a near-baseline transcriptional state, the genes that remain upregulated are related to an alternative-activation transcriptional program. This alterations may aid in maintenance of the Th2 T_RM_ immune environment or be beneficial during subsequent neuroinflammatory challenge by inhibiting pro-inflammatory T cell activation or by enhancing debris clearance.

### Acute *Ts* infection induces gut-to-brain trafficking and transient blood-brain barrier leakiness

We have established that acute *Ts* infection induces rapid and sustained infiltration of leukocytes into the CNS, yet the source of these immune cells remains unclear. In EAE, CD4^+^ T cells are licenced in the intestine and gut-associated lymphoid tissue for subsequent migration into the CNS (83); therefore, we hypothesized that leukocytes may migrate from the intestinal environment to the CNS during *Ts* infection. During acute *Ts* infection, the number of leukocytes and Th2s in the intestinal-draining mesenteric lymph nodes (mLNs) expanded rapidly between 5 and 6 DPI, just prior to their appearance in CNS (**Figure 5A, S5A**). Further, CD4^+^ T cells in the mLN at 5 DPI had increased expression of CD49d, the majority of which were GATA3^+^ Th2s (**Figure 5B-D, S5B**). CD49d is a subunit of the VLA-4 integrin that promotes leukocyte extravasation into the CNS, perhaps indicating increased migration of Th2s into the CNS.

**Figure 5.**
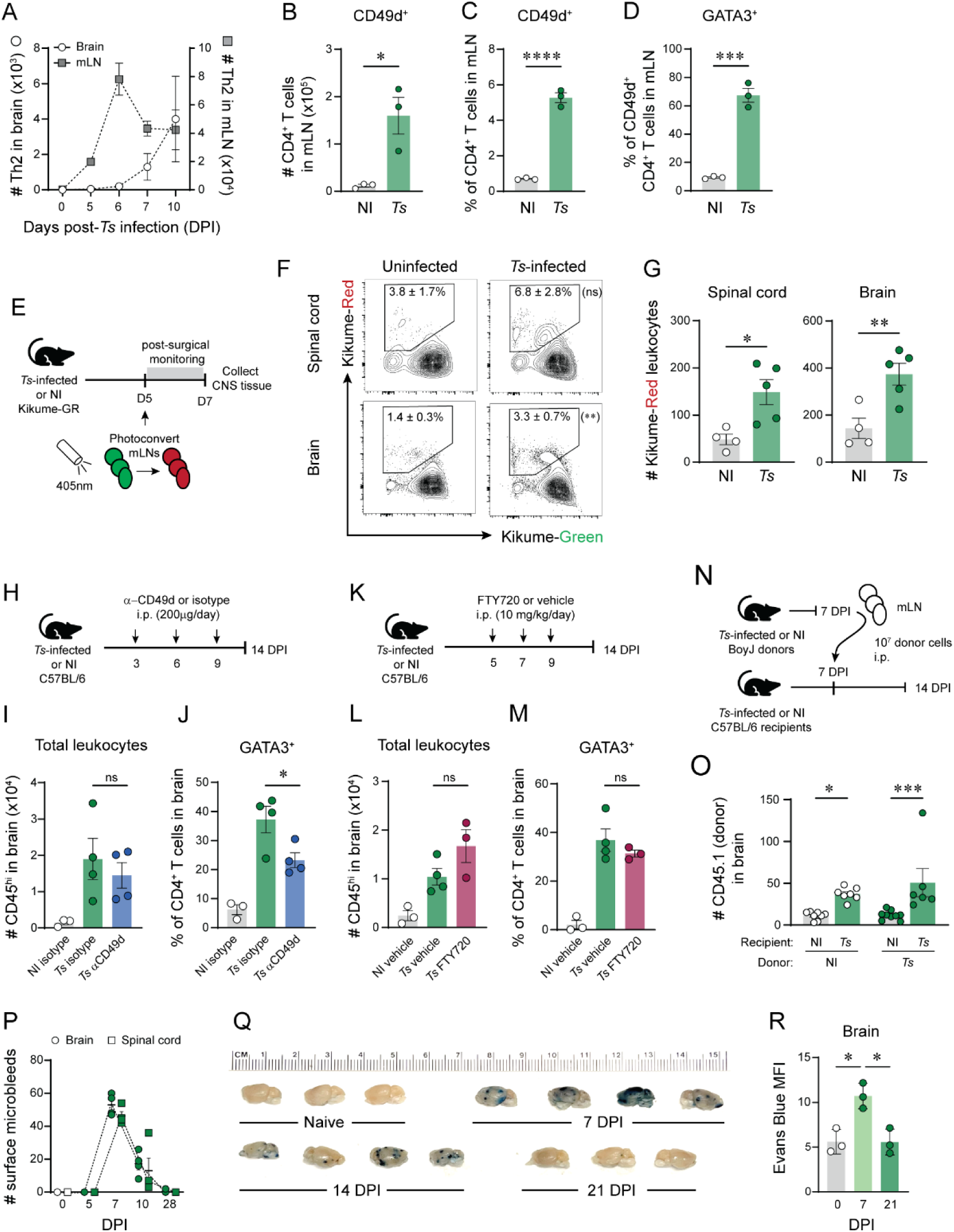
Acute *Ts* infection induces gut-to-brain trafficking and transient BBB leakiness. (A) Number of Th2s in the brain (open circles, left y-axis) and in the mesenteric lymph nodes (mLN) (grey squares, right y-axis) during acute *Ts* infection. (B) Number of CD49d^+^ CD4^+^ T cells, (C) proportion of CD49d^+^ CD4^+^ T cells, and (D) proportion of GATA3^+^ CD49d^+^CD4^+^ T cells in the mLN at 5 DPI. (E) *Ts* or NI KikGR mice underwent abdominal surgery to photoconvert the mLNs at 5 DPI, and CNS tissue evaluated at 7 DPI. (F) Concatenated flow plots of KikGR-Red (photoconverted) vs KikGR-Green CD45^hi^ leukocytes in the spinal cord (top) and brain (bottom) (NI: n=4, *Ts*: n=5). (G) Number of spinal cord- and brain-infiltrating KikGR-Red CD45^hi^ leukocytes. (H) *Ts* or NI mice were treated with 200 μg α−CD49d or isotype control (rat IgG2b) i.p. per day on days 3, 6, and 9 post-*Ts* infection, and CNS infiltration evaluated at 14 DPI. (I) Number of leukocytes (CD45^hi^) in brain. (J) Proportion of GATA3^+^ CD4^+^ T cells in the brain. (K) *Ts* or NI mice were treated with 10 mg/kg FTY720 or vehicle (dH_2_O) i.p. on days 5, 7, and 9 post-*Ts* infection, and CNS infiltration evaluated at 14 DPI. (L) Number of leukocytes (CD45^hi^) in the brain. (M) Proportion of GATA3^+^ CD4^+^ T cells in the brain. (N) MLNs from *Ts* (7 DPI) or NI BoyJ (CD45.1^+^) donors were collected and 10^7^ bulk leukocytes i.p. injected into 7 DPI *Ts-* infected or NI C57BL/6 (CD45.2^+^) recipients, and CNS infiltration evaluated 7 days post-transfer. (O) Number of donor leukocytes (CD45.1^+^) in the brain. (P) Number of microbleeds on the surface of the brain and spinal cord in *Ts-*infected mice. (Q) Photographs of brains from naïve, 7, 14, or 21 DPI *Ts-*infected mice following i.v. injection of Evans Blue dye. (R) Median fluorescent intensity (MFI) of Evans Blue in the brain. Statistics by unpaired t-test (B-D, G), one-way ANOVA with Tukey’s multiple comparisons test (I, J, L, M, R), or two-way ANOVA with Fisher’s LSD test (O, see Figure S5D for ANOVA table).

To test whether immune cells migrated directly from the mLNs to the CNS, we used Kikume Green-Red (KikGR) photoconvertible mice to track cellular migration (84). We performed abdominal surgery at 5 DPI to photoconvert the mLNs of *Ts-*infected or uninfected mice. Mice were recovered for 2 days prior to collection of CNS tissue at 7 DPI to evaluate infiltration of mLN-derived (Kikume-Red^+^) cells (**Figure 5E**). *Ts-*infected mice had more Kikume-Red^+^ leukocytes in the brain and spinal cord compared to uninfected mice (**Figure 5F, G**), suggesting that the CNS-infiltrating cells are at least in-part derived from the mLN. To test the role of CD49d in gut-to-brain trafficking, *Ts-*infected mice were treated with α-CD49d neutralizing antibody (clone PS/2) or isotype control on days 3, 6, and 9 post-*Ts* infection (**Figure 5H**). CD49d neutralization did not affect the total number of CNS-infiltrated leukocytes at 14 DPI, but did reduce the proportion of Th2s in the CNS (**Figure 5I, J**). Due to the increased expression of CD49d on Th2s during acute infection, α-CD49d may have preferentially inhibited migration of Th2s into the CNS. To more broadly inhibit cellular trafficking, *Ts-*infected mice were treated with the S1PR1 agonist FTY720 on days 5, 7, and 9 post-*Ts* infection (**Figure 5K**), which similarly had no effect on total leukocyte nor Th2 infiltration into the CNS, (**Figure 5L, M)**, despite increased retention of activated and polarized CD4^+^ T cells in the mLN (**Figure S5C**). To test an alternative hypothesis that leukocyte migration into the CNS occurs due to other host factors associated with *Ts* infection, we adoptively transferred bulk leukocytes from the mLN of *Ts-*infected (7 DPI) or naïve BoyJ (CD45.1) donors into naïve or *Ts-*infected C57BL/6 (CD45.2) recipients (**Figure 5N**). The ability of transferred donor cells to migrate to the CNS was dependent on the *Ts* infection status of the recipient mouse, regardless of the *Ts* infection status of the donor mouse (**Figure 5O, S5D**). Altogether, these data indicate that *Ts* enhances leukocyte infiltration into the CNS independently of immune priming in the mLN during acute *Ts* infection.

Coinciding with the day post-*Ts* infection that leukocytes can be detected in the CNS (7 DPI), we observed the presence of punctate hemorrhages covering the surface of the brain and spinal cord. The presence of these bleeds was transient and began to subside by 10 DPI, and were completely absent by 28 DPI (**Figure 5P, S5E**). H&E-stained sections of brain tissue similarly revealed the presence of hemorrhages within the CNS parenchyma tissue on 7 and 10 DPI, which resolved by 28 DPI (**Figure S5F)**. The brain had increased permeability to intravenous Evans Blue dye at 7 DPI which returned to baseline by 21 DPI, as measured by the fluorescent signal of Evans Blue and by visual count of the lesion load (**Figure 5Q, R, S5G-H**). These *Ts-*induced bleeds resemble the vascular damage and hemorrhage occasionally described in people with neurological manifestations of Trichinellosis (85-87). The prevailing hypothesis is that circulating *Ts* newborn larvae burrow into the endothelial cells that line the capillaries of the CNS to cause physical disruption of the vasculature; however, there is limited evidence for the presence of *Ts* larvae in the CNS tissue in humans or animal models. Vascular damage could alternatively be induced by inflammatory cascades in response to *Ts* infection, as these bleeds resemble the cerebral microhemorrhages induced upon LPS administration (88-90); however, these mechanisms have yet to be explored.

### Th2s are necessary for recovery from EAE, but not delay in EAE onset

These data establish that *Ts* infection potently modifies the CNS immune landscape, but the effect of these changes on EAE outcomes in *Ts-*infected mice remain unclear. We hypothesized that Th2 and/or Tregs in the periphery or resident in the CNS may contribute to the delay of EAE onset and the recovery of EAE symptoms. To evaluate the necessity of the type 2 immune response, STAT6-deficient mice were infected with *Ts* prior to immunization with MOG_35-55_/CFA. Due to their inability to effectively clear the adult-stage *Ts* infection (91), *Stat6*^-/-^ mice did not tolerate the 400 L1 dose used throughout this study, and developed more severe and sustained weight loss, a higher muscle-stage L1 burden, and a failure of immune cells to expand in the mLN (**Figure S6A-C**). Therefore, a lower dose of 100 *Ts* L1 was used to evaluate the effect of STAT6 deficiency on EAE outcomes, which promoted EAE amelioration similar to the high-dose infection in wildtype mice (**Figure S6D-F**). As expected, *Stat6*^-/-^ CD4^+^ T cells had impaired expansion of Th2s in the periphery and the CNS, but an enhanced Treg population in the brain during acute *Ts* infection (**Figure S6G, H**).

*Ts-*infected *Stat6*^-/-^ mice had a similar delay in EAE onset as S*tat6*^+/-^ littermate controls (**Figure 6A**). During the priming phase of EAE, RORγt expression was reduced in *Ts/*EAE SLOs in a STAT6-independent manner (**Figure 6B**, **S6I**), and IL-17A expression by *Ts/*EAE CD4^+^ T cells was similarly reduced following either PMA or MOG_35-55_ stimulation independently of STAT6 (**Figure 6C, S6J**). This reduced Th17 polarization mirrored a *Ts*-mediated but STAT6-independent increase in Tregs, but was de-coupled from STAT6-dependent polyclonal IL-13 expression in *Ts/*EAE mice (**Figure 6C, S6K**). These data suggest that type 2 immunity is not necessary for *Ts-*mediated inhibition of Th17 polarization, nor the delay in EAE onset.

**Figure 6.**
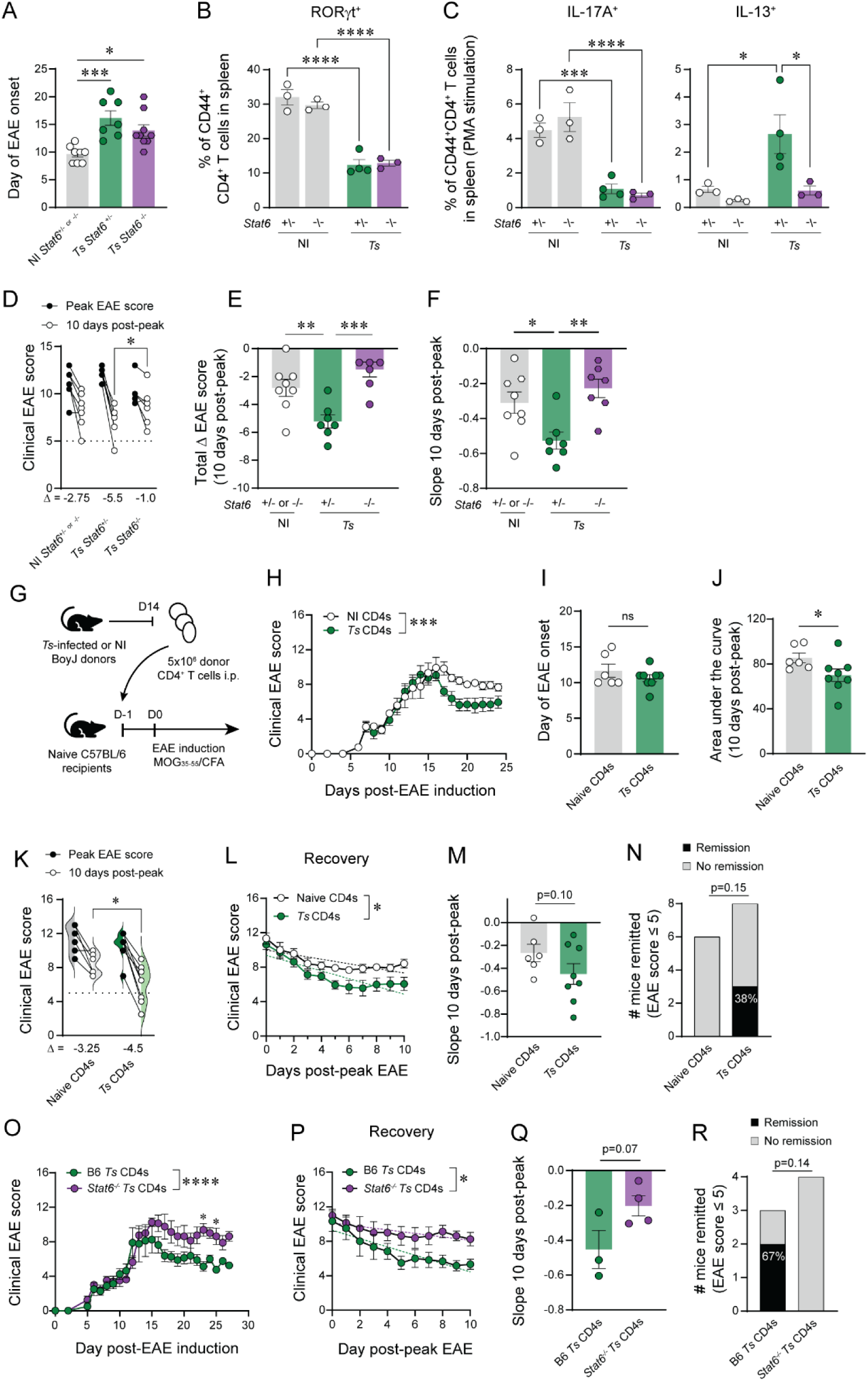
Th2s are necessary for recovery from EAE, but not delay in EAE onset. (A-F) *Ts-* infected (100 *Ts* L1 o.p.) or NI *Stat6^-/-^* or *Stat6^+/-^*littermates were immunized with MOG_35-55_/CFA at 28 DPI. Clinical data pooled from 2 independent experiments. (A) Day of EAE onset. (B) Proportion of RORγt^+^ CD44^hi^CD4^+^ T cells in the spleen. (C) Proportion of IL-17A^+^ (left) or IL-13^+^ (right) CD44^hi^CD4^+^ T cells in the spleen following PMA/ionomycin stimulation. (D) Paired analysis of the peak EAE score vs 10 days post-peak. Delta (Δ) = median change in EAE score. (E) Change in EAE score (Δ) in the 10-day post-peak interval. (F) Slope of the linear regression of the 10-day post-peak interval for individual mice. (G-N) CD4^+^ T cells were isolated from the spleens and mLNs of *Ts-*infected or NI BoyJ (CD45.1) donors at 14 DPI, and 5×10^6^ CD4^+^ T cells were transferred i.p. into naïve C57BL/6 recipients 1 day prior to MOG_35-55_ immunization. Data pooled from 2 independent experiments. (H) Clinical EAE scores of recipients of NI or *Ts* CD4^+^ T cells. (I) Day of EAE onset. (J) Area under the curve in the 10 days post-peak disease. (K) Paired analysis of the peak EAE score reached vs EAE score 10 days post-peak. Delta (Δ) = median change in EAE score. (L) EAE scores of 10 days post-peak EAE for individual mice. Dashed lines show linear regression. (M) Slope of the linear regression of the 10-day post-peak interval for individual mice. (N) Number of mice remitted (EAE score ≤ 5). (O-R) CD4^+^ T cells were isolated from the spleens and mLNs of *Ts-*infected C57BL/6 or *Stat6^-/-^* (CD45.2) donors at 14 DPI, and 5×10^6^ CD4^+^ T cells were transferred i.p. into naïve BoyJ (CD45.1) recipients 1 day prior to MOG_35-55_ immunization. (O) Clinical EAE scores of mice that received CD4^+^ T cell transfers from *Ts-*infected C57BL/6 or *Stat6^-/-^* donors. (P) EAE scores of 10 days post-peak EAE for individual mice. Dashed lines show linear regression. (Q) Slope of the linear regression of the 10-day post-peak interval for individual mice. (R) Number of mice remitted (EAE score ≤ 5). Statistics by one-way ANOVA with Tukey’s multiple comparisons test (A, E, F), two-way ANOVA with Tukey’s (B, C) or Sidak’s multiple comparisons test (D, H, K, O), unpaired t-test (I, J, M, Q), ANCOVA test for equivalency of linear regressions (L, P), or one-sided Fisher’s exact test (N, R).

Although *Stat6*^-/-^ mice had delayed EAE onset, they did not resolve their clinical EAE scores like STAT6-sufficient *Ts/*EAE mice (**Figure 6D, S6L-M**). The magnitude of change in EAE score during the 10-day post-peak period and the rate of clinical EAE recovery were significantly reduced in *Ts-*infected *Stat6*^-/-^ mice compared to *Stat6^+/-^* littermates (**Figure 6E, F)**. These data suggest that infiltration of type 2 immune cells into the CNS is necessary to prime the neuroinflammatory environment to promote recovery and repair upon subsequent neuroinflammatory challenge. STAT6-mediated remission could be mediated through a variety of CNS-infiltrated type 2 immune cells, including Th2s, ILC2s, or alternatively activated macrophages, as well as other STAT6-expresing CNS-resident cells such as microglia. During acute *Ts* infection, microglial activation was exacerbated (**Figure S6N**), but whether STAT6-deficient microglia polarize in such a manner that could sustain ongoing T cell inflammation, rather than promote resolution or repair, remains unknown.

To assess the sufficiency of *Ts-*primed CD4^+^ T cells, rather than other type 2 immune cells, to mediate amelioration of EAE, we adoptively transferred CD4^+^ T cells from NI or acutely *Ts-* infected (14 DPI) congenic donor mice (BoyJ) into uninfected C57BL/6 recipients 1 day prior to MOG_35-55_ immunization (**Figure 6G**). As expected, *Ts*-infected donor cells had increased expression of CD44, FOXP3, and GATA3 compared to uninfected donors (data not shown). The infection status of the donor did not affect the day of EAE onset (**Figure 6H-I**), however, mice that received *Ts-*primed CD4^+^ T cells recovered from severe EAE better than those that received cells from uninfected donors, including significantly lower total disease burden, lower EAE scores at endpoint, greater change in EAE score following peak disease, and more mice achieved clinical remission (EAE score ≤5) (**Figure 6J-N**). To confirm that recovery is driven by Th2s, rather than other CD4^+^ T cell populations, transfer of donor CD4^+^ T cells from STAT6-deficient donors was unable to promote clinical recovery (**Figure 6O-R**). These data highlight that exogenous supply of *Ts-*primed Th2s is sufficient to induce recovery during neuroinflammatory challenge, and that clinical benefit can be derived from *Ts-*mediated immune modulation independently of overt infection.

### Therapeutic *Ts* infection promotes clinical remission

For *Ts*-mediated neuroimmune modulation to be beneficial as immunotherapy, *Ts* infection must be able to inhibit an already-inflamed environment. To this end, we investigated the therapeutic potential of *Ts* infection in the C57BL/6 x SJL/J (B6xSJL) F1 relapsing-remitting EAE model. First, we verified that the B6xSJL F1 strain has ameliorated EAE symptoms when prophylactically infected with *Ts* 4 weeks prior to MOG_35-55_ immunization, and indeed *Ts*-infected B6xSJL F1 mice had slightly delayed EAE onset and underwent fewer clinical relapses (ΔEAE score >4, reaching at least an EAE score of 7) (**Figure S6O-R**). To evaluate the efficacy of *Ts* infection as a therapeutic, B6xSJL F1s were immunized with MOG_35-55_/CFA, then infected with *Ts* after resolution of the first wave of EAE symptoms (day 18 post-EAE induction). Mice that remained uninfected developed subsequent clinical relapses, whereas mice that were therapeutically infected with *Ts* remained in clinical remission and continued to recover body weight (**Figure 7A-D**). Therapeutic *Ts* infection promoted infiltration of Th2s into the CNS and reduced the numbers and proportions of Th17s and Tregs ‒ reflective of the CD4^+^ T cell phenotypes seen during prophylactic *Ts*/EAE (**Figure 7E-F**). These data highlight the ability of *Ts* to shift the inflammatory tone of the CNS even after an initial inflammatory event has occurred. Altogether, the immunomodulatory nature of *Ts* provide a promising novel therapeutic strategy to prevent or treat neuroinflammatory disease.

**Figure 7.**
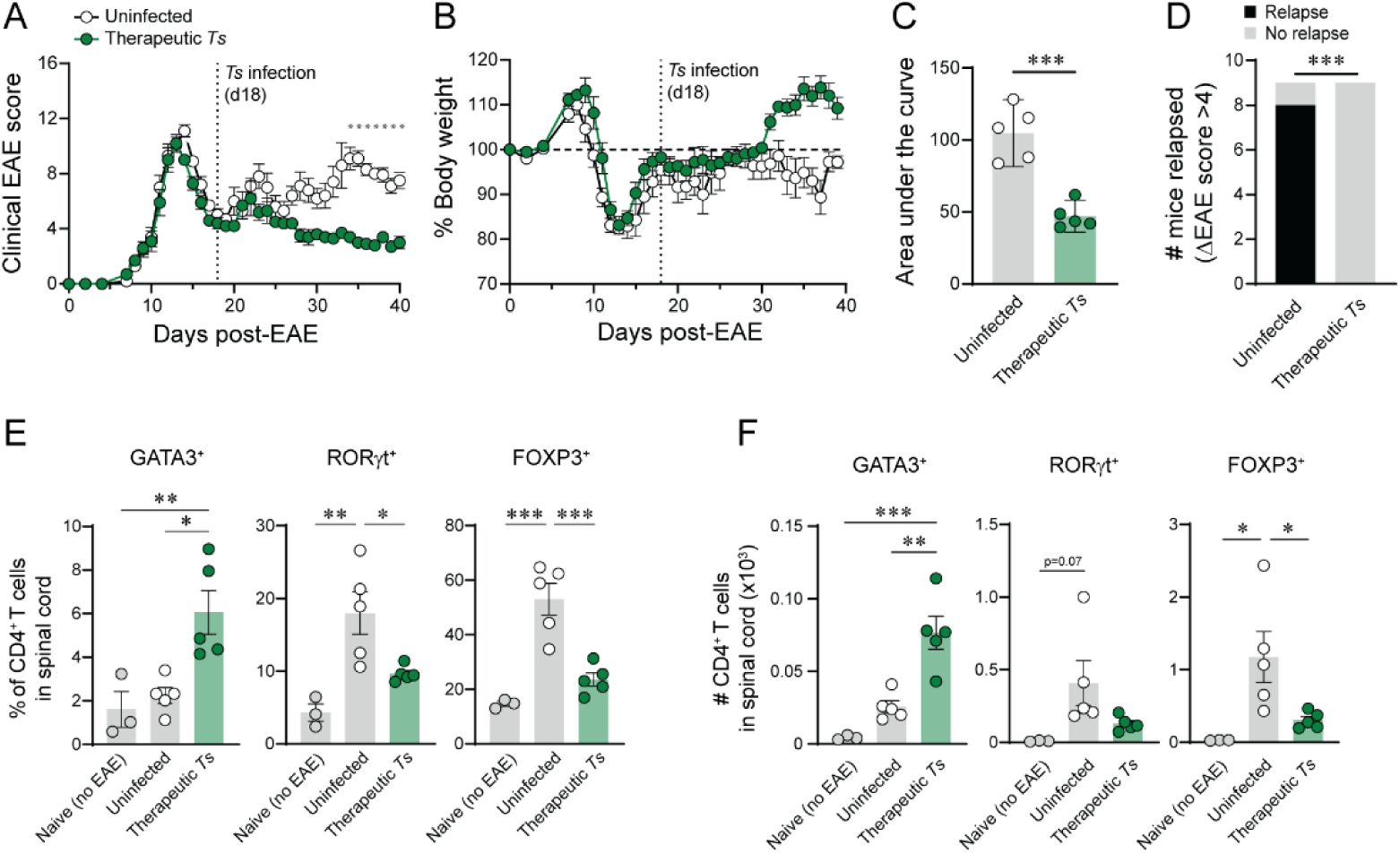
Therapeutic *Ts* infection promotes EAE remission. C57BL/6J×SJL F1 mice were immunized with MOG_35-55_/CFA, then infected with 400 *Ts* L1 after resolution of the first wave of EAE symptoms. (A) Clinical EAE scores. and (B) percent body weight of EAE mice before and after *Ts* infection. (C) Area under the curve from the day of *Ts* infection to endpoint. (D) Number of mice relapsed (Δ EAE score > 4). Data pooled from 2 independent experiments. (E) Proportion and (F) number of GATA3^+^, RORγt^+^, and FOXP3^+^ CD4^+^ T cells in the spinal cord on day 40 post-EAE. Statistics by 2-way ANOVA with Sidak’s multiple comparisons test (A), unpaired t-test (C), one-sided Fisher’s exact test (D), one-way ANOVA with Tukey’s multiple comparisons test (E, F).

## Discussion

The presence of type 2 immune cells and their effector molecules in the CNS or meningeal borders have increasingly-appreciated functions in promoting CNS health, including through regulation of neuronal circuit development (92-94), resolution of spinal cord and neuronal injury (95, 96), and improvements in cognitive function at homeostasis and during age-related or Alzheimer’s Disease-related cognitive decline (97-101). In the context of EAE and MS, the presence of IL-4 during T cell priming promoted IL-10 production by encephalitogenic Th1s (102), CNS-intrinsic IL-4 signaling was required for alternative activation of microglia to restrain inflammation (103), and therapeutic administration of IL-4 into the CNS promoted EAE recovery via IL-4Ra signaling in neurons (104); altogether suggesting that CNS-intrinsic type 2 immune signaling could have therapeutic value in MS and other neurodegenerative or neurotraumatic disease contexts.

Here, we provide data bridging previous observations that helminth infections can ameliorate EAE via peripheral immunomodulation with the emerging understanding of the role of type 2 immune activity within the CNS. In addition to a STAT6-independent delay in EAE onset that is consistent with previously described outcomes in *Ts*-infected rat models of EAE (105, 106), we demonstrate that *Ts* elicits a population of CNS-resident Th2 T_RM_ cells that are associated with neuroprotective effects, including rapid recovery from EAE-induced paralysis and reduced pathogenic inflammation and demyelination in the CNS. Importantly, the protective effects of *Ts* infection were apparent both prophylactically and therapeutically, suggesting that *Ts*-mediated activation of type 2 immunity can promote translationally relevant effects that could be harnessed for neuroprotection or repair.

We initially hypothesized that the delayed EAE onset in *Ts*-infected mice could be mediated by direct Th2 inhibition of Th1 and/or Th17 effector function, or by alternative macrophage activation, which have been shown to ameliorate EAE (107-110). However, the *Ts*-elicited delay in EAE onset was STAT6-independent, suggesting an alternative mechanism to type 2 immune-mediated inhibition. One possibility is that the delay is mediated by a regulatory cell, such as Tregs or regulatory B cells (Bregs). As demonstrated here and by others (106, 111), Tregs are potently activated following *Ts* infection. However, amelioration of EAE by naturally-occurring or engineered Tregs is almost exclusively characterized by a reduction in clinical severity, rather than delayed onset (47, 48, 112, 113). *Ts-*infected EAE mice also have fewer Tregs at peak EAE than NI controls, which could reflect reduced generation of Tregs in response to damage, as opposed to Treg-mediated resolution of inflammation. Further, *Ts*-infected STAT6-deficient mice had a compensatory increase in Tregs following acute infection, yet displayed a similar delay in EAE onset, suggesting that the delay may be Treg-independent. We have not evaluated regulatory B cell populations in this study, however, an increase in Bregs has been noted in people with MS with naturally-acquired helminth infections (114, 115) and in *Ts*-infected mice (116). Bregs are critical for restraining EAE initiation, whereby pan-B cell or regulatory B cell depletion prior to EAE induction have been shown to increase EAE severity and time to onset (117, 118). Bregs have further been associated with EAE recovery (119, 120). Alternatively, tolerogenic DCs have been demonstrated to reduce EAE severity following *in vitro* polarization (121, 122), including stimulation with *Ts-*derived products (123, 124).

Tolerogenic DC differentiation is a STAT6-independent process (125), suggesting that this may contribute to the diminished activation of pro-inflammatory Th17 and MOG_35-55_-reactive CD4^+^ T cells and delayed EAE-induced autoimmune neuroinflammation in *Ts*-infected animals. During the immune priming phase of EAE, we detected an approximately equal number of MOG-IA^b+^ Tetramer^+^ CD4^+^ T cells, with a reduction in the frequency and number of Th17s. However, it remains unclear if there is a proportional increase in a different effector cell type, as we did not detect a substantial increase in MOG-reactive Tregs or Th2s. One possibility is the expansion of lineage-negative anergic T cells, characterized by expression of FR4 and CD73, similar to those that have been described in the context of oral food tolerance and suppression of autoimmunity (126-129). Supporting this possibility, we identified a significant upregulation of *Nt5e* (CD73) in the SLOs of *Ts/*EAE SLOs, which may reflect an accumulation in “anergic” antigen-specific cells. Further, T helper lineage-negative cells may be predominantly activated by cDC2s (128), the classical type 2 immune DC subtype, and thus could pose another potential mechanism by which *Ts-*primed DCs contribute to ameliorated EAE.

Although other helminths have been investigated for EAE-modifying potential with generally positive results (28), the clinical and immunological phenotypes we describe following *Ts* infection provide several advantages over other helminths for translational potential, including long-lasting protection from EAE onset, perhaps due to the persistent nature of the *Ts* natural lifecycle, along with potent alterations to the CNS immune landscape. Due to limited investigation of the CNS neuroimmune landscape in other models of HIT in EAE, it remains unknown whether the *Ts*-mediated alterations to the neuroimmune landscape described here are unique to *Ts* infection, including transient blood-brain barrier permeability, leukocyte influx, and maintenance of a Th2 T_RM_ populations. Subsequent evaluation of each of these phenotypes in isolation will untangle which are responsible for the recovery from EAE – a clinical phenotype unique to *Ts* among other models of HIT in EAE.

It is important to emphasize that pre-clinical investigation of *Ts* in modulation of EAE will provide important mechanistic understanding of *Ts-*mediated type 2 protective immunity in the CNS, rather than suggest that *Ts* should be directly used as immunotherapy. Although *Ts* may have played a historically significant role in educating the human immune system, public health efforts to eradicate *Trichinella* species from domestic food supplies have drastically reduced the burden of Trichinellosis for a net benefit on human health. By studying mechanisms by which this human-relevant helminth modulates the CNS neuro-immune landscape, we may be able to identify translationally relevant pathways of *Ts-*mediated neuroprotection, including generation of CNS-resident Th2 T_RM_ cells, while avoiding damaging aspects of infection, such as *Ts-* mediated hemorrhage. Altogether this study highlights the previously unappreciated neuro-modulatory role of *Ts* infection, which can be harnessed for preventative or therapeutic benefit in neuroinflammatory disease.

## Resource Availability

This study did not generate new unique reagents. RNA sequencing and Nanostring nCounter data will be deposited at the NCBI GEO repository following peer review. Any additional information required to reanalyze the data reported in this paper is available from the lead contact upon request.

## Acknowledgements

The authors acknowledge support from MS Canada (DRG-1032809 to LCO), the Canadian Institutes for Health Research (PJT-186265 to LCO), the Research Corporation for Science Advancement Scialog award (28637 to LCO and UBE), the National Institutes of Health (NIH RO1 AI151599, RO1AI169770, RO1AI180644 to MCS), the Rutgers 2024 Busch Biomedical Grant Program (AS), and the MS Canada endMS Personnel Award program (NMF, SJP, TW). This work was supported by Life Sciences Institute ubcFLOW core and the UBC Centre for Disease Modelling supported by the UBC GREx Biological Resilience Initiative, the computational resources and services provided by Advanced Research Computing at UBC, UBC SBME-Seq, the UBC Centre for Heart Lung Innovation Molecular Phenotyping Core, and the NIH tetramer core. We are grateful to Annie Cierna for her technical insights.

## Methods

### Mouse strains and husbandry

Mice were housed at the University of British Columbia (UBC) under specific pathogen-free conditions at the Center for Disease Modeling. C57BL/6J (000664), BoyJ (B6.SJL-PtprcaPepcb/BoyJ, 002014), CAG-KikGR (Tg(CAG-KikGR)33Hadj/J, 013753) (84), and STAT6-knockout (B6.129S2(C)-Stat6tm1Gru/J, 005977) (130) mice were ordered from The Jackson Laboratories and bred in-house. SJL (SJL/J-CrHsd, 052) mice were obtained from Envigo Labs and bred in-house prior to breeding SJL/J males with C57BL/6 females to generate C57BL/6×SJL (B6×SJL) F1 mice.

Experimental mice were all female and age-matched within each experiment between 6-12 weeks of age for all experiments. If ordered in from The Jackson Laboratories, 5–8-week-old mice were randomized within 48 hours of delivery to minimize cage-specific effects and allowed to acclimatize for at least one week before any experimental procedures were performed. Mice were randomly assigned into experimental groups prior to the first treatment or intervention. Up to 5 mice per cage were housed in ventilated Ehret cages prepared with BetaChip bedding and had ad libitum access to irradiated PicoLab diet 5053 and reverse osmosis/chlorinated (2–3 ppm)-purified water. Housing rooms were maintained on a 14/10-hour light/dark cycle with temperate and humidity ranges of 20–22°C and 40–70%, respectively. All experiments were performed according to guidelines from UBC Animal Care Committee and Biosafety Committee-approved protocols.

For Evans Blue permeability assay, mice were maintained and bred in a pathogen-free, virus Ab-free facility at Rutgers New Jersey Medical School Comparative Medicine Resources. Healthy, 7–12-week-old mice were selected for experimental groups from purchased or bred colonies, without using specific randomization or blinding methods. These studies were reviewed and approved by the Institutional Animal Care and Use Committee at Rutgers-the State University of New Jersey. All experiments were conducted according to the principles specified in the Guide for the Care and Use of Laboratory Animals, Institute of Animal Resources, National Research Council, Department of Health, Education and Welfare (US National Institutes of Health).

### *Trichinella spiralis* (*Ts*) infections and quantification

*Ts* stocks were maintained by serial *in vivo* passaging in immune-sufficient mice. First-stage larvae (L1) were recovered from infected muscle tissue by digestion in pepsin-HCl (1% each) for 3-5 hours at 37°C, shaking at 180 rpm. Larvae were washed by settling through several changes of sterile PBS, allowing for 5-minute settling time between washes. L1s were counted under a light microscope and resuspended to the desired concentration in sterile PBS. Mice were infected by oral gavage with a total of 400 L1, or 100 L1 where specified, in 200 μL PBS. Quantification of muscle-encysted L1 was performed by digesting a pre-weighed tissue sample as described above, then counted using a light microscope and normalized for an L1 burden per gram of tissue.

### Experimental autoimmune encephalomyelitis (EAE) inductions and clinical monitoring

EAE was induced by immunization with 200 μg of MOG_35-55_ peptide (MEVGWYRSPFSRVVHLYRNGK; GenScript) emulsified in CFA (400 μg desiccated *Mycobacterium tuberculosis* H37 Ra mixed with incomplete Freund’s adjuvant (BD Biosciences)). The emulsion was injected subcutaneously in a total volume of 100 μL in the hind flank under isoflurane anaesthesia. Intraperitoneal (i.p.) injections of 200 ng pertussis toxin (List Biological Laboratories) were administered on the day of immunization and 48 hours later.

Mice were evaluated for clinical health and EAE severity daily after the onset of EAE symptoms. EAE severity was scored according to a composite 16-point scale, which measures mobility impairments in each limb and the tail, as previously described (131, 132). Each limb is graded from 0 (asymptomatic) to 3 (complete paralysis), the tail is graded from 0 (asymptomatic) to 2 (limp tail), and the righting reflex is scored from 0 to 2, with 0 assigned for normal righting reflex, 1 for slow righting reflex, and a 2 for a delay of more than 5 seconds in the righting reflex. Each criterion was measured in 0.5-point increments, thus composite scores range from 0 (completely asymptomatic) to 16 (fully quadriplegic with limp tail and significantly delayed righting reflex). Only mice that survived until dedicated experimental endpoints were included in area under the curve calculations, and only mice that developed EAE were included in clinical curves or quantifications, apart from incidence curves in which all immunized mice were included.

### Antibody and drug treatments

FTY720 (Selleckchem, S5002) was administered at 10 mg/kg in dH_2_0 by i.p. injection. FTY720 solution or dH_2_O vehicle was injected on days 5, 7, and 9 post-*Ts* infection. Anti-CD49d (clone PS/2, UBC AbLab) or rat IgG2b isotype control was administered in three doses of 200 μg each on days 3, 6, and 9 post-*Ts* infection.

### Photoconversion surgery

Photoconvertible mice (CAG-KikGR) were infected with 400 *Ts* L1s by oral gavage or left uninfected. 5 days post-infection, all mice underwent abdominal surgery to photoconvert the mesenteric lymph node complex. Mice were anaesthetized to a surgical plane with isoflurane, then administered 20 mL/kg Lactated Ringer’s Solution, 5 mg/kg meloxicam, and 0.1 mg/kg buprenorphine subcutaneously. The abdomen was shaved, the surgical site sterilized with hibitane and ethanol, and 8 mg/kg bupivacaine was injected subcutaneously along the planned incision site. A 1-2 cm incision was made down the midline of the abdomen, and a slightly smaller incision along the underlying muscle layer. The cecum was gently exteriorized and the mLNs located, then the surrounding tissue was shielded with aluminum foil. The mLNs were photoconverted by exposure with a 405 nm 160 mW/cm^2^ laser positioned 15 cm away for 60 seconds. The organs were returned to the abdominal cavity, then incisions were closed with 5-0 Monocryl sutures and sealed with GLUture. Lactated Ringer’s Solution and buprenorphine were re-administered as needed every 8 hours, and meloxicam was administered every 24 hours. Mice were euthanized 48 hours post-surgery.

### Histology and image quantification

Mice were euthanized by CO_2_ inhalation and transcardially perfused with at least 20 mL cold PBS. Brain and SCs were submerged in 10% neutral buffered formalin for 24 hours then transferred to 70% ethanol for storage at 4°C prior to paraffin embedding. Spinal cords were cut into 4 approximately equal-length sections for cross-sectional tissue slices. Paraffin blocks were sectioned at a thickness of 5-7 μm and serial sections were stained with hematoxylin and eosin (H&E) and Luxol Fast Blue (LFB) for cellular infiltration and demyelination, respectively. Paraffin embedding, sectioning, and staining were completed by a commercial histology service (Wax-it Histology Services Inc.).

Slides were imaged with a Zeiss Axio Observer Z1 and an AxioCam 105 microscope camera. Brightfield images were acquired with a 10X objective. 3 sections per spinal cord were scored for infiltration in a blinded manner. The median score between 3 independent evaluations was used for quantification. Inflammation/infiltration scores were assigned using a 4-point system as follows: 0, no infiltration; 1, mild increase in cellularity/infiltration in perivascular space; 2, several separate regions of perivascular infiltration/cuffing; 3, severe perivascular infiltration and 1-2 regions of parenchymal infiltration; 4, severe diffuse infiltration with several parenchymal clusters. LFB-stained slides were acquired at 10X magnification with consistent white balancing and exposure times within individual experiments. Myelination was quantified in 3 sections per spinal cord using ImageJ Fiji processing software. White matter regions of interest (ROI) were manually selected, then colour thresholding (hue, saturation, brightness) was determined based on naïve sections and applied to each ROI. Myelination was calculated as a percent area of the ROI.

For Evans Blue dye blood-brain barrier permeability assay, mice were injected i.v. with 2% Evans Blue dye 1 hour prior to euthanasia. Mice were transcardially perfused with 20 mL ice-cold PBS followed by 4% PFA. Brains were fixed overnight in 4% PFA at 4°C, then cryoprotected in 30% sucrose for 24-48 hours, then frozen in OCT. 20 μm sagittal sections were cryo-sectioned and mounted, then stained with DAPI (1:2000 dilution for 20 minutes). Coverslips were mounted with ProLong Gold. Slides were imaged using a Keyence BZ-X710 inverted fluorescence microscope at 4X resolution with DAPI channel exposure of 1/5 s and TexasRed Channel exposure (within Evans Blue excitation range) of 1.2 s. Using Fiji processing software, the average RGB pixel values were taken, and the average red value quantified for each pixel.

### Leukocyte recovery, single cell suspension processing, and ex vivo stimulation

Single cell suspensions from spleens and lymph nodes were prepared by mechanical dissociation and passage through a 70 μm nylon mesh. Spleen suspensions were incubated with ACK lysis buffer for 5 minutes on ice to remove red blood cells. To isolate leukocytes from CNS tissue, mice were transcardially perfused with at least 20 mL cold PBS, then brain and spinal cord tissues were passed through a 70 μm nylon mesh filter followed by resuspension in room temperature 40% isotonic Percoll and centrifugation at 500 × g for 15 minutes with no brake. Cell pellets were resuspended in FACS buffer (PBS supplemented with 2% NCS and 2-5 mM EDTA) or in stimulation media for intracellular staining for cytokine responses. For isolation of dural meninges (adapted from (133)), skull caps were collected and stored in RPMI + 2% NCS. Under a dissection microscope, meninges were peeled from the edges of the skull cap using sharp forceps. The meninges were incubated at 37°C in a shaking incubator at 180 rpm for 30 minutes in 1 mL digestion buffer containing 1 mg/mL collagenase D (Roche, catalog #11088858001) and 50 μg/mL DNase I (Sigma, catalog #EN0521) in RPMI. Digested meninges were filtered through a 70 μm cell strainer and washed with 2% NCS-supplemented PBS.

For polyclonal cytokine responses, single cell suspensions were stimulated with 0.1 μg/mL PMA (Sigma) and 0.1 μg/mL ionomycin (Sigma) in the presence of Brefeldin A (eBioscience/ThermoFisher) and GolgiSTOP (BD Biosciences) for 5 hours at 37°C. All data points represent whole organ suspensions from individual mice. For antigen-specific cytokine responses, single cell suspension were stimulated in RPMI-based complete tissue culture media (10% FBS, sodium pyruvate, MEM non-essential amino acids, HEPES, penicillin/streptomycin) containing 50 μg/mL MOG_35-55_ for 5 hours in the presence of Brefeldin A and GolgiSTOP (spleen, lymph nodes), or with 20 μg/mL MOG_35-55_ overnight (∼16 hours) and in the presence of Golgi inhibitors for the final 5 hours of stimulation (brain, spinal cord), or with 20 μg/mL *Ts* L1 antigen for 72 hours and in the presence of Golgi inhibitors for the final 5 hours of stimulation (spleen).

*Ts* L1 antigen for stimulation was generated by collecting *Ts* L1s as described above using 0.2 μm filter-sterilized digest buffer and sterile PBS, then homogenizing for 10 minutes at 30 Hz in an autoclaved 2 mL round-bottom tube containing ∼50 μL of glass beads (425–600 μm). The resulting homogenate was centrifuged at 12 000 × g for 10 minutes, then the protein content in the supernatant quantified by bicinchoninic acid assay (BCA) as per kit manufacturer’s instructions (Pierce BCA protein kit, ThermoFisher catalog #23225).

### CD4^+^ T cell enrichment

CD4^+^ T cells were negatively isolated using a cocktail of biotinylated antibodies (2.5 μg/mL each CD11b (M1/70), CD11c (N418) F4/80 (BM8), B220 (RA3-6B2), CD19 (1D3), and CD8a (53-6.7)). Splenocytes were isolated as described above and pooled with mLN-derived cell suspensions at 10^8^ cells/mL in isolation buffer (2% NCS, 1 mM EDTA). Cells were Fc-blocked using 50 μL/mL rat serum, then incubated with 20X antibody cocktail for 15 minutes at 4°C. Streptavidin-bound magnetic beads (StemCell Technologies, #50001) were added (50 μL/mL) and incubated for 3 minutes at room temperature, then another 3-minute incubation in a magnet, and the flowthrough used for downstream applications. Purity was evaluated by flow cytometry of pre- and post-enrichment cell suspension.

### Flow cytometry

Cells were washed with cold PBS prior to staining with Fixable Aqua viability dye (Invitrogen/ThermoFisher) for 15 minutes at 4°C in the dark. Subsequently, cells were washed with FACS buffer and incubated with 2 μg/mL α-mFcγR (clone 24G2, AbLab) for 5-10 minutes at RT. If tetramer staining was performed, cells were incubated with 7 μg/mL I-Ab PE-MOG (GWYRSPFSRVVH) tetramer or irrelevant peptide (human CLIP) control (PVSKMRMATPLLMWQA) at room temperature in the dark for 1 hour, then washed with FACS buffer prior to extracellular antibody staining. Tetramers were pre-conjugated and tetramerized by the NIH Tetramer Core Facility. Pre-determined concentrations of fluorophore-labelled antibodies were added for a final volume of 100 μL of Fc-block and antibodies in FACS buffer. Wells were mixed thoroughly then incubated for 30 minutes at 4°C in the dark. If intracellular staining was performed, cells were next incubated with FOXP3/Transcription Factor Fixation/Permeabilization buffer (eBioscience/ThermoFisher) for at least 30 minutes (up to overnight) at 4°C in the dark, followed by 2 washes with permeabilization buffer. Fluorophore-labelled antibodies for intracellular targets were diluted in 100 μL permeabilization buffer per well and incubated with cells for 40 minutes at room temperature in the dark. For intracellular MBP detection, purified anti-MBP was stained with other intracellular antibodies as described above, followed by a 20-minute incubation at room temperature with the secondary antibody.

Samples were run on a CytoFLEX LX N3-V5-B3-Y5-R3-I0 (BD Biosciences) flow cytometer. Analysis was performed using FlowJo v10.7 (BD Life Sciences). Unless otherwise stated, representative flow cytometry plots depict concatenated files from a given experimental group.

### Microglia isolation and cell sorting for RNA sequencing

For microglia isolation prior to RNA extraction, mice were perfused with 30 mL cold HBSS (no calcium, no magnesium) supplemented with actinomycin D (5 μg/mL) and triptolide (10 μM) and stored in the same perfusion buffer on ice until processing. Brains were cut into small pieces with a scalpel and placed in 5mL HBSS containing actinomycin D (5 μg/mL), triptolide (10 μM), and anisomycin (27.1 μg/mL) (HBSS + inhibitors). Once all samples were processed, 50 μg/mL Papain (diluted 1:1 with HBSS) and 66 μg/mL DNAse1 were added then samples were incubated in a 37°C water bath for 15 minutes. Samples were triturated with a P1000 pipette tip incubated for another 15 minutes, repeated once more for a total of 45 minutes of incubation. Samples were then filtered through a pre-soaked 70 μm filter and centrifuged at 400 × g at 4°C for 10 minutes. The pellet was resuspended in 5.6 mL HBSS + inhibitors and added to a 15 mL conical tube containing 2.4 mL 90% isotonic Percoll (2.133 mL Percoll + 0.266 mL HBSS per sample) and inverted to mix, then 2 mL HBSS was overlaid to create a discontinuous gradient. Samples were centrifuged at 400 × g for 20 minutes at 4°C with no break or acceleration. The supernatant was aspirated to remove the myelin layer and all but the bottom ∼400 μL of buffer and the cell pellet. The pellet was washed in FACS buffer then stained for FACS. Microglia were FACS-purified based on viability staining, then identified by being TCRβ/B220 negative, CD45.2^mid^CD11b^+^ and P2RY12^+^. Samples were run on a CytoFLEX SRT cell sorter (BD Biosciences). Samples were sorted into 50% FBS-containing HBSS + inhibitors and temporarily stored on ice. Immediately following isolation, samples were centrifuged and resuspended in 1 mL TRIzol for RNA extraction. A maximum of 8 mice were processed per day.

### RNA isolation, sequencing, and gene expression analysis

For TRIzol RNA extractions, cell pellets were resuspended in 1 mL TRIzol (Invitrogen) and stored at -70°C (to a maximum of 3 months) prior to RNA extraction. RNA was extracted as per manufacturer’s instructions. Briefly, samples were brought to room temperature and incubated for 5 minutes. 0.2 mL chloroform was added and incubated for 2 minutes. Samples were centrifuged for 15 minutes at 12 000 × g at 4°C. Upper aqueous phase was removed and placed in a new tube, and the chloroform extraction and centrifugation were repeated once. 10 μg of RNase-free glycogen (ThermoFisher) was added to aid in visualization of the pellet during subsequent precipitation steps. 0.5 mL of isopropanol was added to the aqueous phase, incubated for 10 minutes, then centrifuged for 10 minutes at 12 000 × g at 4°C. The supernatant was removed and the pellet was resuspended in 1 mL ice-cold 75% ethanol. Samples were vortexed then centrifuged for 5 minutes at 7500 × g at 4°C. The ethanol wash was repeated twice. The pellet was air-dried for 10 minutes then resuspended in 20 μL dH2O and stored at - 70°C prior to sequencing.

Sample quality control was performed using the Agilent 2100 Bioanalyzer for purity and quantity, and only samples with an RNA integrity score (RIN) >8 were processed for library preparation following the standard protocol for Illumina Stranded mRNA prep (Illumina), and sequencing as performed on the Illumina NextSeq2000 with paired End 59bp x59 bp reads. Library preparation and sequencing were performed by the UBC SBME Sequencing (UBC SBME-Seq) core facility. FASTQ files were mapped and aligned using Salmon v1.10.1 (134). Reads were mapped to an index created using the GRCm38 mouse transcriptome assembly with decoy sequences generated using SalmonTools.

Differential gene expression analysis was performed in RStudio on mapped and aligned file outputs from Salmon. The tximeta package was used to convert transcript quantifications to Ensembl mouse transcriptome (GRCm39) gene counts (135). Batch correction was performed with the limma package (136). Briefly, a linear model was fit to the data, and the component associated with the batch was removed for subsequent plotting and data exploration. DESeq2 was used for variance stabilization, PCA, and differential gene expression analysis (137). Criteria for significance throughout RNA analyses were: log_2_ fold change > or < 2, p_adj_ < 0.05, and base mean > 50. Heatmaps were generated with the pheatmap function with samples organized by group, and genes were hierarchically clustered using Euclidean distance by Ward’s method. Heatmap data show mean-normalized batch-corrected expression values. PERMANOVA (999 permutations) was performed using pairwise adonis (138). The ClusterProfiler package was used for gene set enrichment analysis (GSEA) of all mouse GO terms and published RNA-seq datasets, as well as calculations of q-values and normalized enrichment scores (139). Relevant microglial gene sets were identified from the MGEnrichment App curated mouse database (140). Gene count values were extracted following log_2_ transformation with variance stabilizing transformations and batch correction. All p-values were adjusted for multiple comparisons using FDR cutoff of 0.05 (5%).

For Nanostring nCounter analysis, a portion of the spleen or the iLN ipsilateral to the immunization injection site were collected and stored in RNAlater (Thermo) at -20°C. RNA was extracted using the PureLink RNA Mini Kit (Invitrogen) as per manufacturer’s instructions. RNA was analyzed for RNA concentration and purity using an Agilent 22100 BioAnalyzer. 100 ng of RNA per sample was run on the Nanostring nCounter Mouse Host Response panel. Analysis was performed using ROSALIND (Rosalind.bio). Each sample passed hybridization quality controls, and transcripts were removed with numbers of reads below the background threshold determined by negative controls. Data was normalized to positive controls and to housekeeping genes based on the ROSALIND geNorm algorithm. Probe counts were determined by dividing counts within a lane by the geometric mean of the normalizer probes in the same lane. Tissues (inguinal lymph node, spleen) were analyzed separately due to differences in baseline expression levels and differences in housekeeping gene stability. Gene set enrichment analysis was performed in R with ClusterProfiler on select mouse GO terms (139, 141).

### Statistical analysis

Statistical analyses were performed using GraphPad Prism (GraphPad Software, Version 10.4). Data was assessed for normal distribution by Shapiro-Wilk test. Two-tailed nonparametric Mann-Whitney test, two-tailed t-test, one-way ANOVA with Tukey’s multiple comparisons test, Kruskal-Wallis with Dunn’s multiple comparisons test, 2-way ANOVA with Tukey’s or Šídák’s multiple comparisons test, multiple Mann-Whitney tests with Benjamini, Krieger, and Yekutieli multiple comparisons test (FDR=1%), Log-rank (Mantel-Cox) test, or Fisher’s exact test were used as specified in figure legends. Statistical analysis for RNA sequencing data was performed in RStudio, and Nanostring nCounter data analyzed in ROSALIND. All data points from a given experiment were included unless otherwise stated to be removed from clinical EAE curves (i.e., asymptomatic mice) or that an ROUC outlier test (Q=1%) was performed. Unless otherwise stated, data shown is one representative experiment of at least 2 independent repeats. Data are shown as mean ± SEM unless otherwise stated. Symbols: ns p > 0.05, ∗ p < 0.05; ∗∗ p < 0.01; ∗∗∗ p < 0.001; ∗∗∗∗ p < 0.0001.

## Supplemental Information

**Figure S1.**
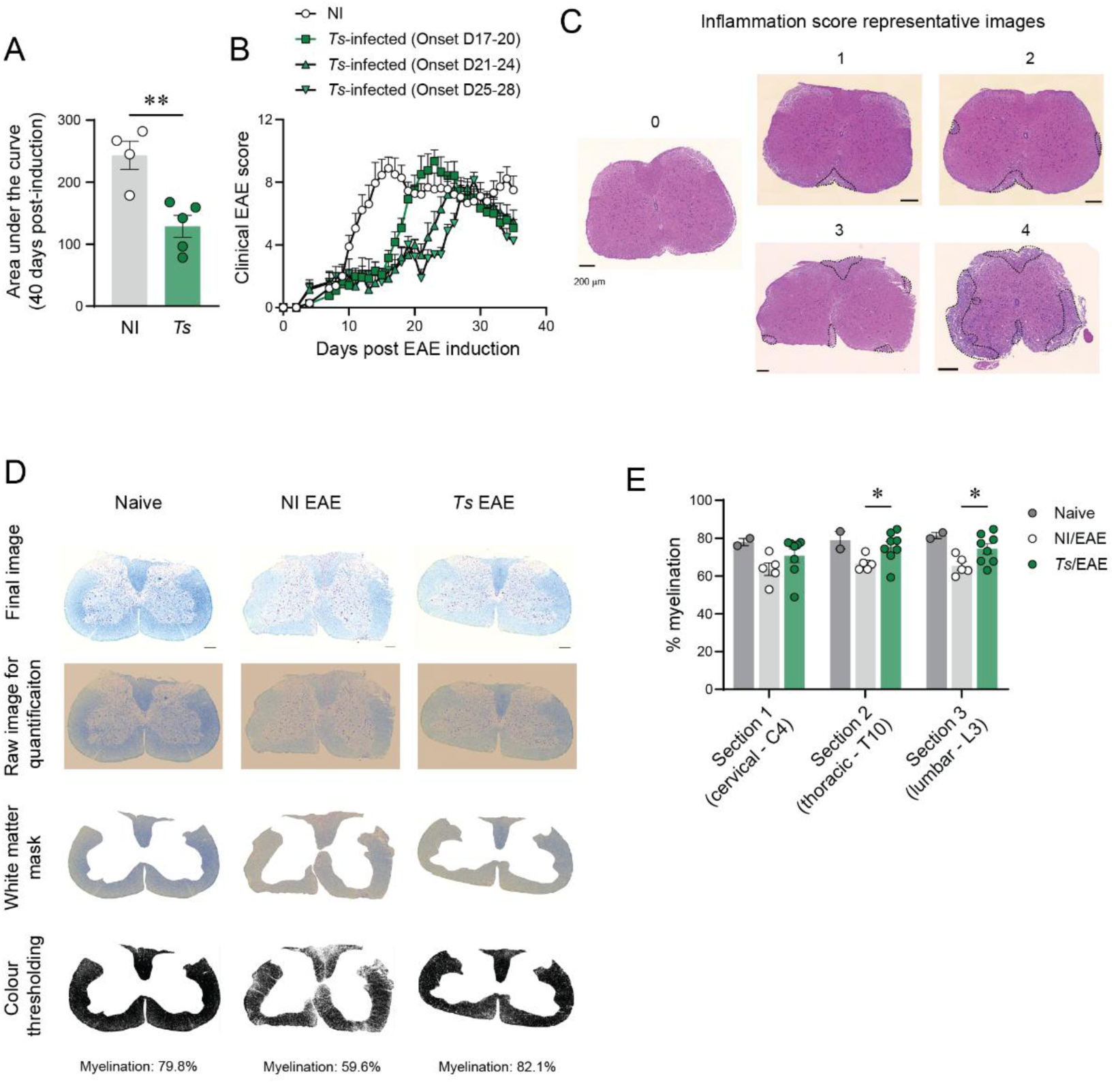
Chronic *Ts* infection delays EAE onset and promotes recovery. (A) Area under the curve for the full 40-day course of EAE from one representative experiment (B) Clinical EAE scores from (Figure 1D), with *Ts/*EAE mice stratified by day of EAE onset (day 17-20, day 21-24, or day 25-28). (C) Representative hematoxylin and eosin (H&E) staining of spinal cord sections for inflammation scores 0-4. Areas of cellular infiltration used to categorize samples are outlined in black dashed lines. Scale bar = 200μm. (D) Representative workflow for quantification of myelination in Luxol Fast Blue (LFB) stained sections. Briefly, images were acquired using identical exposure time and white balancing, the white matter manually masked and the colour thresholds adjusted based on values for a naïve control from same experiment, then %myelination quantified per white matter area. (E) Myelination quantification separated by approximate spinal cord section.Statistics by unpaired t-tests (A, E).

**Figure S2.**
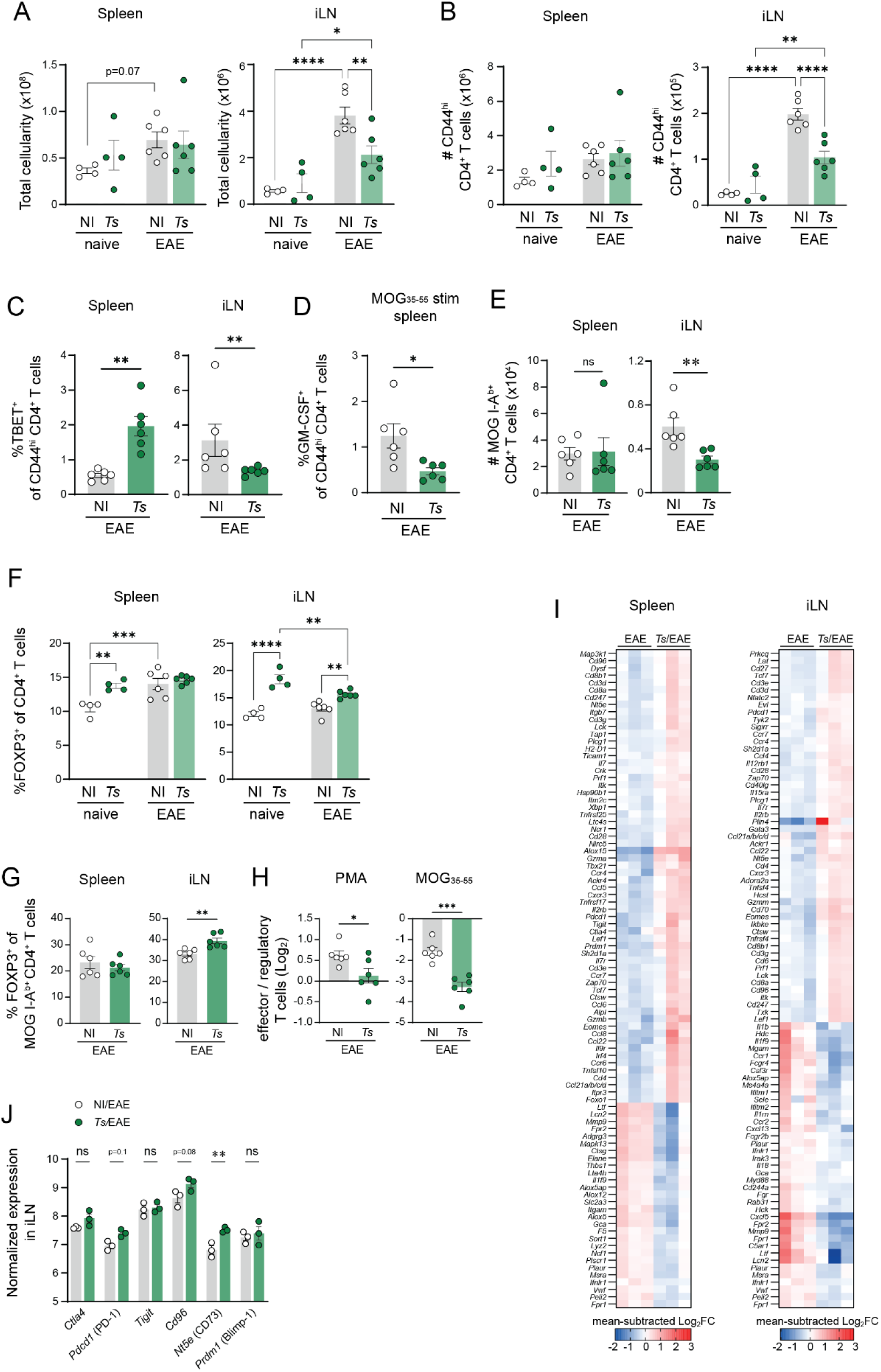
*Ts* alters immune priming in EAE. Spleens and inguinal lymph nodes (iLNs) were harvested on day 9 post-EAE (or non-EAE naïve controls) from *Ts-*infected or not infected (NI) mice. (A) Total cellularity by hemocytometer counts in the spleen and iLN following preparation of single cell suspensions. (B) Number of CD44^hi^ CD4^+^ T cells in the spleen and iLN. (C) Proportion of TBET^+^ CD44^hi^CD4^+^ T cells in the spleen and iLN. (D) Proportion of CD44^hi^CD4^+^ T cells that express GM-CSF in the spleen and iLN following MOG_35-55_ stimulation. (E) Number of MOG-IA^b+^ CD4^+^ T cells n the spleen and iLN. (F) Proportion of FOXP3^+^ CD4^+^ T cells in the spleen and iLN. (G) Proportion of FOXP3^+^ MOG-IA^b+^ CD4^+^ T cells n the spleen and iLN. (H) Log2-transformed ratio of the number of effector T cells (producing IL-17A^+^ and/or IFNγ^+^) to the number of FOXP3^+^ Tregs in the spleen following PMA/ionomycin or MOG_35-55_ stimulation. (I) Nanostring nCounter differential gene expression summary in the spleen and iLN. Genes are hierarchically-clustered and shown as mean-subtracted Log_2_ Fold Change within each tissue.(J) Normalized expression in the spleen and iLN of select co-inhibitory molecules.Statistics by two-way ANOVA with Tukey’s (A, B, F) or Sidak’s multiple comparisons test (J), unpaired t-test (C-E, G, H), or a Generalized Linear Model for Nanostring differential gene expression (performed in Rosalind) (I).

**Figure S3.**
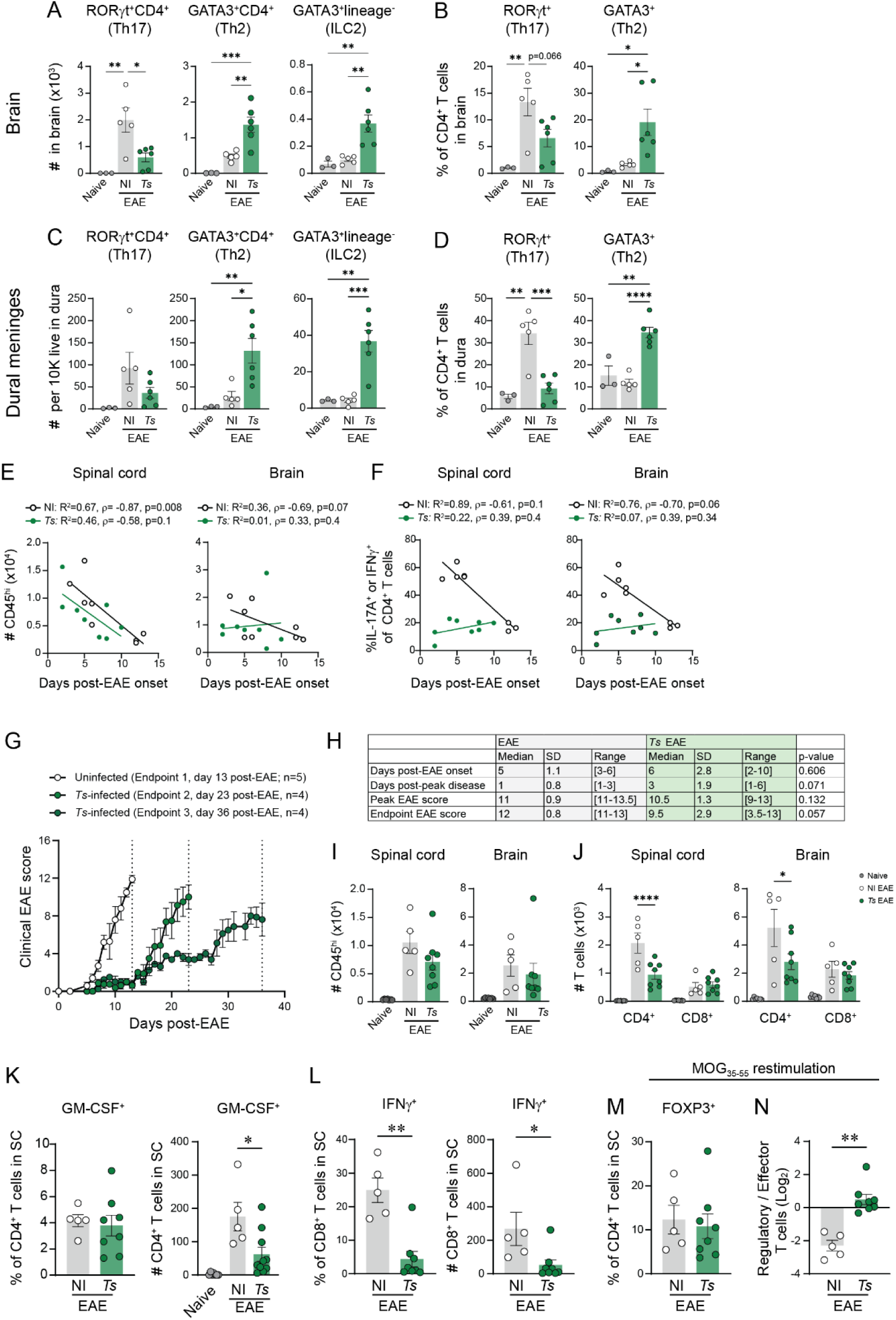
Chronic *Ts* infection alters T cell and microglia responses during peak EAE. (A) Number of brain-infiltrating RORγt^+^CD4^+^ T cells (Th17), GATA3^+^CD4^+^ T cells (Th2) and GATA3^+^lineage^-^(TCRβ/B220/CD11b/CD11c^-^) CD45^hi^ cells in the brain on day 40 post-EAE induction. (B) Proportion of RORγt^+^ or GATA3^+^ CD4^+^ T cells in the brain. (C) Number of Th17, Th2, and ILC2 in the dural meninges per 10 000 live cells. (D) Proportion of RORγt^+^ or GATA3^+^ CD4^+^ cells in the dural meninges. (E) Correlation between number of CD45^hi^ cells in the spinal cord or brain and the number of days post-EAE onset at experimental endpoint. (F) Correlation between the proportion of IFNγ and/or IL-17A-expressing CD4^+^ T cells in the spinal cord, brain, or spleen, and the number of days post-EAE onset at experimental endpoint. (G) Clinical EAE scores of mice used CNS cytokine evaluation at peak EAE. (H) Clinical EAE metadata for NI EAE or *Ts* EAE mice euthanized at peak EAE. (I) Number of CD45^hi^ leukocytes in the spinal cord and brain at peak disease. (J) Number of CD4^+^ and CD8^+^ T cells in the spinal cord and brain at peak disease. (K) Proportion and number of GM-CSF^+^ CD4^+^ T cells in the spinal cord following PMA stimulation. (L) Proportion and number of IFNγ^+^ CD8^+^ T cells in the spinal cord. (M) Proportion of FOXP3^+^ CD4^+^ Tregs in the spinal cord, and (N) the ratio of Tregs to effector (IFNγ^+^ and/or IL-17A^+^) CD4^+^ T cells in the spinal cord following overnight MOG_35-55_ stimulation. Statistics by one-way ANOVA with Tukey’s multiple comparisons test (A-D), linear regression and Spearman correlation (E-F), two-way ANOVA with Tukey’s multiple comparisons test (J), Mann-Whitney tests (K-N).

**Figure S4.**
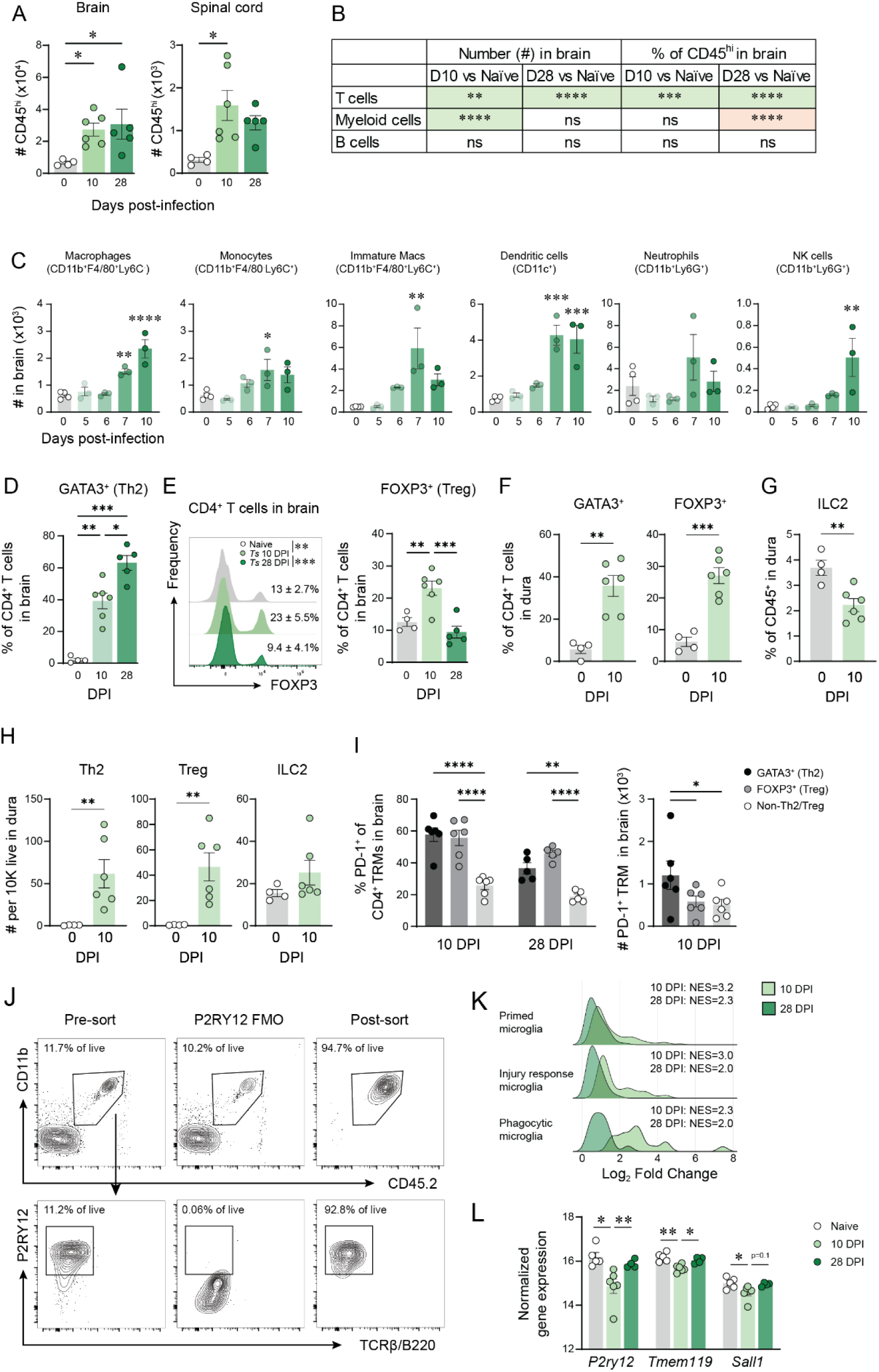
Acute *Ts* infection drives remodeling of the neuroimmune landscape. (A) Number on CD45^hi^ leukocytes in the brain or spinal cord of naïve or *Ts-*infected mice at 10 DPI or 28 DPI. (B) Two-way ANOVA table of leukocyte populations in the brain (related to Figure 4A). Green colour shows increase in *Ts* vs NI, orange shows decrease in *Ts* vs NI. (C) Number of various myeloid cell populations in the brain during acute *Ts* infection. (D) Proportion of GATA3^+^ Th2s of CD4^+^ T cells in the brain at 0, 10, or 28 DPI. (E) Concatenated histograms (n=4-6) of FOXP3 expression in CD4^+^ T cells in the brain (left). Inset data shows %FOXP3^+^ of CD4^+^ T cells ± SD. Quantification of proportion of FOXP3^+^ CD4^+^ T cells in the brain (right). (F) Proportion of GATA3^+^ and FOXP3^+^ CD4^+^ T cells, and (G) proportion of ILC2s of (GATA3^+^lineage^-^(TCRβ/B220/CD11b/CD11c^-^) CD45^hi^ cells) of total CD45^+^ in the dural meninges at 10 DPI. (H) Number of Th2, Treg, and ILC2 in the dural meninges per 10 000 live cells. (I) Proportion (left) and number (right) of GATA3^+^, FOXP3^+^, or non-Th2/Treg CD4^+^ T_RM_ (CD69^+^) that express PD-1. (J) Representative flow plots of gating strategy of microglia FACS for RNA sequencing, depicting microglia pre-FACS purification (left), the P2RY12 FMO used to identify microglia (centre), and post-sort microglia (right). (K) Gene set enrichment analysis ridgeplots of 10 DPI and 28 DPI microglia RNAseq compared to 3 published microglia gene sets. (L) Normalized gene expression of select microglia homeostatic markers by RNA sequencing in naïve, 10 DPI, and 28 DPI microglia. Statistics by one-way ANOVA with Tukey’s multiple comparisons test (A, D, E, I, L), or two-way ANOVA with Sidak’s multiple comparisons test (C), or unpaired t-test (F, G, H).

**Figure S5.**
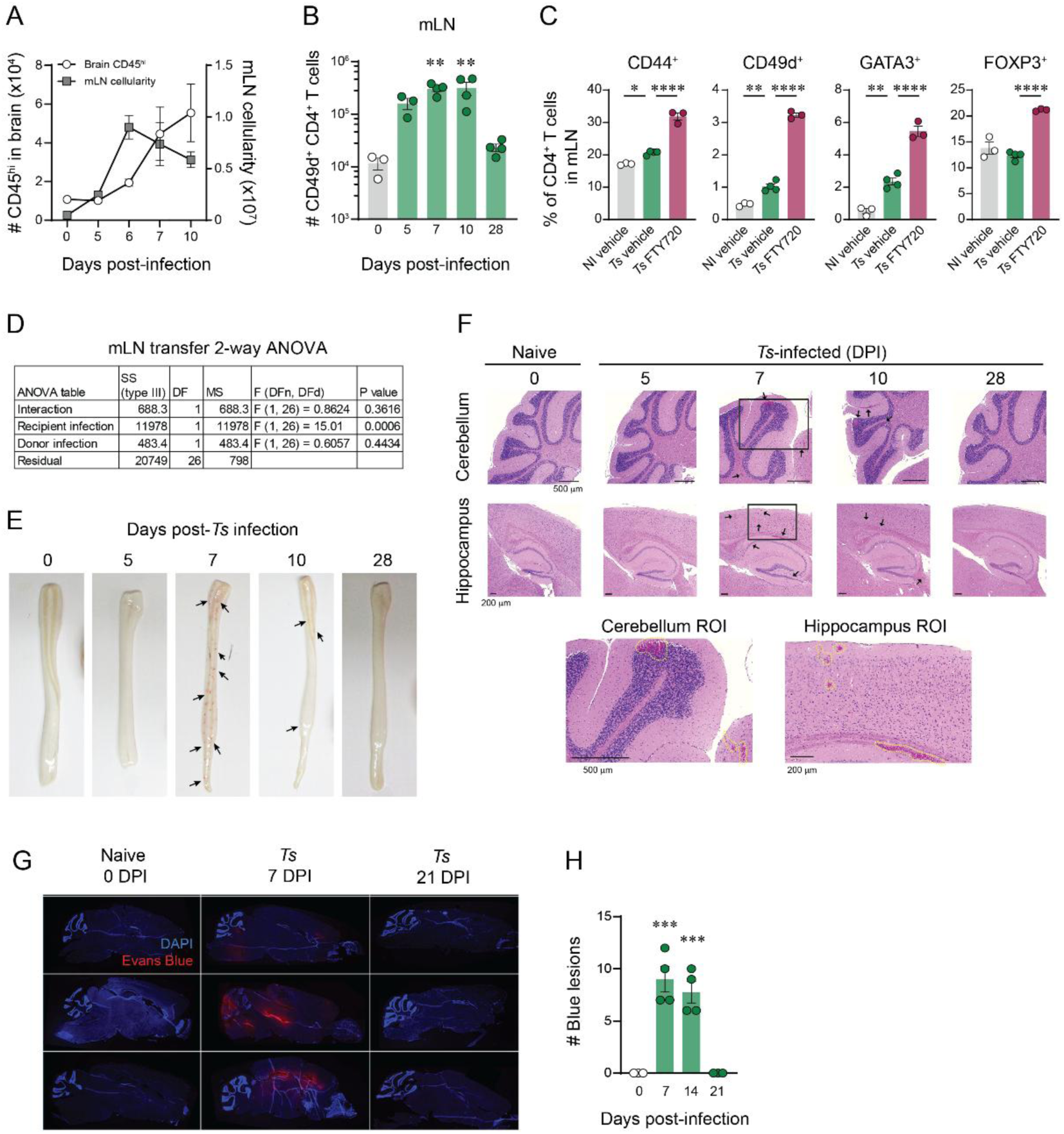
Acute *Ts* infection induces gut-to-brain trafficking and transient BBB leakage. (A) Number of CD45^hi^ leukocytes in the brain (open circles, left y-axis) and total mLN cellularity (grey squares, right y-axis). (B) Number of CD49d^+^CD4^+^ T cells in the mLN during acute *Ts* infection. (C) Proportion of CD44^+^, CD49d^+^, GATA3^+^, or FOXP3^+^ CD4^+^ T cells in the mLN at 14 DPI of FTY720- or vehicle-treated *Ts* or NI mice. (D) ANOVA table of data shown in Figure 5O. (E) Representative photographs of perfused spinal cords at various days post-*Ts* infection. (F) Representative sagittal sections of brain cerebellum (top) or hippocampus (bottom) of NI or *Ts-* infected mice on various days post-infection. Scale bar = 500 μm (cerebellum) or 200 μm (hippocampus). Region of interest (ROI) selected from 7 DPI samples. (G) Immunofluorescent images of brains from NI or *Ts-*infected mice injected i.v. with 2% Evans Blue dye 1 hour prior to fix-perfusion. (H) Quantification of Evans Blue lesions. Statistics by one-way ANOVA with Dunnet’s multiple comparisons test (each DPI vs NI) (B, H) or Tukey’s multiple comparisons test (C), two-way ANOVA (D).

**Figure S6.**
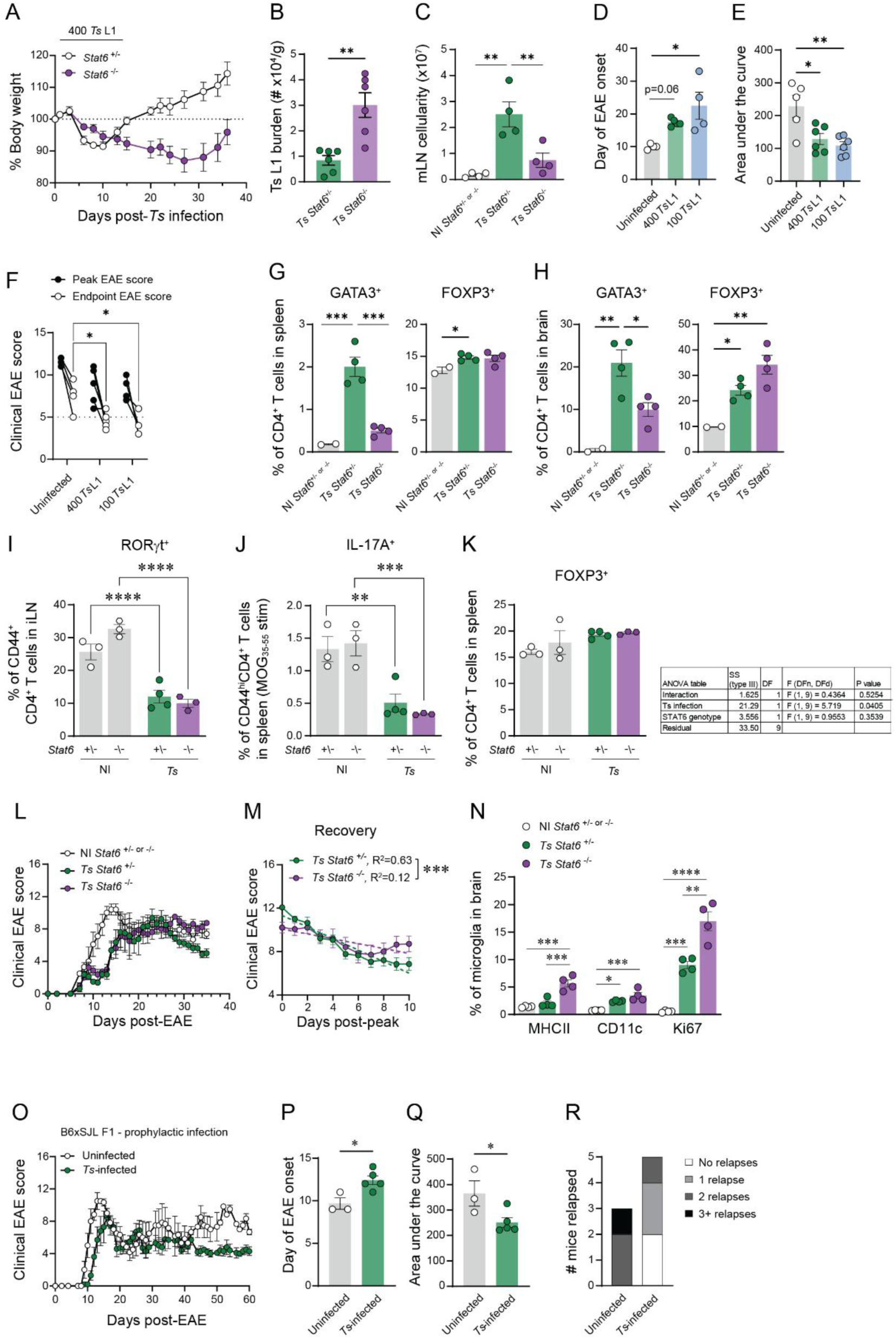
Th2s are necessary for recovery from EAE, but not delay in EAE onset. *Stat6^+/-^* or *Stat6^-/-^* littermates were infected with 400 *Ts* L1 (A-C). (A) Percent body weight during infection. (B) *Ts* L1 burden in the muscle tissue at 28 DPI. (C) Hemocytometer counts of mLN cellularity at 14 DPI. (D-F) C57BL/6 mice were infected with 100 or 400 *Ts* L1 or left not infected (NI) 4 weeks prior to MOG_35-55_ immunization. (D) Day of EAE onset. (E) Area under the curve over 40 days of EAE. (F) Paired analysis of the peak EAE score reached vs EAE score at experimental endpoint. (G-H) *Stat6^+/-^* or *Stat6^-/-^* littermates were infected with 100 *Ts* L1 or left NI and evaluated at 10 DPI. Proportional expression of GATA3^+^ or FOXP3^+^ CD4^+^ T cells in the spleen (G) or brain (H). (I-K) *Stat6^+/-^* or *Stat6^-/-^* littermates were infected with 100 *Ts* L1 or left NI 4 weeks prior to MOG_35-55_ immunization and evaluated at day 9 post-EAE induction. (I) Proportion of RORγt-expressing CD44^+^CD4^+^ T cells in the iLN. (J) Proportion of IL-17A-expressing CD44^+^CD4^+^ T cells in the spleen following MOG_35-55_ stimulation. (K) Proportion of FOXP3-expressing CD4^+^ T cells in the spleen and associated ANOVA table. (L-M) CD4^+^ T cells were isolated from the spleens and mLNs of *Ts-*infected or NI BoyJ (CD45.1) donors at 14 DPI, and 5×10^6^ CD4^+^ T cells were transferred i.p. into naïve C57BL/6 recipients 1 day prior to MOG_35-55_ immunization. (L) Clinical EAE scores of recipient mice. (M) EAE scores for 10 days post-peak disease for individual mice. Dashed lines depict linear regression. (N) Proportion of microglia in the brain of *Ts-*infected *Stat6^+/-^* or *Stat6^-/-^* mice at 10 DPI that express MHCII, CD11c, or Ki67. (O-R) C57BL/6xSJL F1 mice were infected with 400 *Ts* L1 4 weeks prior to MOG_35-55_/CFA EAE induction. (O) Clinical EAE scores. (P) Day of EAE onset. (Q) Area under the curve for full 60-day course of EAE. (R) Number of mice that experienced 0-3+ clinical relapses. Statistics by unpaired t-test (B, P, Q), one-way ANOVA with Tukey’s multiple comparisons test (C-E, G, H, N), two-way ANOVA with Tukey’s multiple comparisons test (F, I, J, K), or Sidak’s multiple comparisons test and ANCOVA test for equivalency of linear regressions (M).

| Day of EAE | EAE only 1 | EAE only 2 | EAE only 3 | EAE only 4 | EAE only 5 | EAE only 6 | EAE only 7 | EAE only 8 | EAE only 9 | EAE only 10 | EAE only 11 | EAE only 12 | EAE only 13 |
| --- | --- | --- | --- | --- | --- | --- | --- | --- | --- | --- | --- | --- | --- |
| 0 | 0 | 0 | 0 | 0 | 0 | 0 | 0 | 0 | 0 | 0 | 0 | 0 | 0 |
| 2 | 0 | 0 | 0 | 0 | 0 | 0 | 0 | 0 | 0 | 0 | 0 | 0 | 0 |
| 4 | 0 | 0 | 0 | 0 | 0 | 1 | 0 | 0 | 3.5 | 1.5 | 4 | 4 | 0 |
| 7 | 0 | 1 | 2 | 1 | 0 | 3 | 0 | 0 | 4.5 | 4 | 4 | 4.5 | 4.5 |
| 8 | 1 | 0 | 1.5 | 0 | 0 | 3 | 1.5 | 1.5 | 4.5 | 4 | 4.5 | 4.5 | 5.5 |
| 9 | 0 | 2 | 2.5 | 4 | 0 | 0 | 1.5 | 3 | 7.5 | 5 | 4 | 4.5 | 4.5 |
| 10 | 4 | 8 | 1.5 | 4.5 | 1.5 | 1.5 | 4 | 4.5 | 9 | 5.5 | 3.5 | 4 | 5.5 |
| 11 | 4 | 8.5 | 4 | 8 | 1.5 | 1.5 | 4 | 9 | 12 | 5 | 5.5 | 3.5 | 4.5 |
| 12 | 5.5 | 8.5 | 4.5 | 10 | 3.5 | 2.5 | 5 | 11 | 11 | 5.5 | 5 | 5.5 | 5 |
| 13 | 8.5 | 10 | 6.5 | 10 | 1 | 2 | 5.5 | 11 | 12 | 5 | 6.5 | 7 | 7 |
| 14 | 11 | 11 | 11.5 | 10 | 2 | 2.5 | 8.5 | 11 | 11 | 4 | 8 | 9 | 7.5 |
| 15 | 11 | 11.5 | 10.5 | 10 | 5 | 4.5 | 9 | 10.5 | 9 | 4 | 12 | 12 | 7.5 |
| 16 | 11 | 11.5 | 10 | 9 | 6.5 | 7 | 9 | 10.5 | 9 | 5.5 | 12 | 11 | 10 |
| 17 | 12 | 12 | 9 | 9 | 4 | 5.5 | 12 | 8 | 9 | 5 | 10 | 11 | 11 |
| 18 | 11 | 10 | 7.5 | 7.5 | 5 | 7 | 12 | 9 | 9 | 5.5 | 11 | 10 | 11 |
| 19 | 8.5 | 7.5 | 7.5 | 6.5 | 5 | 6.5 | 10.5 | 9 | 8.5 | 6 | 11 | 8 | 10 |
| 20 | 7 | 7.5 | 6.5 | 7 | 7.5 | 4.5 | 9 | 9 | 7 | 7.5 | 10 | 7.5 | 10 |
| 21 | 7 | 7.5 | 7 | 8 | 5.5 | 4 | 9 | 11 | 8.5 | 6 | 9 | 8.5 | 9.5 |
| 22 | 7 | 9 | 5.5 | 11 | 4.5 | 4.5 | 9 | 11 | 8.5 | 7.5 | 8 | 9 | 8 |
| 23 | 10 | 8 | 5 | 9 | 3.5 | 3 | 11 | 11 | 8 | 7.5 | 9.5 | 11 | 9 |
| 24 | 9.5 | 8 | 4.5 | 8 | 5 | 5.5 | 10 | 11 | 8 | 7 | 9 | 10.5 | 10 |
| 25 | 10 | 8.5 | 4 | 8 | 5 | 4.5 | 11 | 10 |  | 7 | 9 | 9 | 10 |
| 26 | 9 | 9 | 4 | 6.5 | 7 | 5 | 9 | 10 |  | 5.5 | 9 | 10 | 8.5 |
| 27 | 8 | 7.5 | 4.5 | 6 | 8 | 4 | 9 | 9.5 |  | 7 | 9 | 11 | 9 |
| 28 | 8 | 7 | 5 | 5 | 6.5 | 5 | 9 | 9 |  | 5.5 |  |  | 9 |
| 29 | 8 | 8.5 | 4.5 | 5.5 | 6.5 | 4 | 9 | 9 |  | 6 |  |  |  |
| 30 | 8.5 | 10.5 | 3.5 | 5.5 | 7.5 | 4 | 10 | 9 |  | 5.5 |  |  |  |
| 31 | 6.5 | 8 | 4.5 | 7.5 | 7.5 | 4 | 9 | 9 |  | 5 |  |  |  |
| 32 | 8 | 9 | 5 | 10.5 | 7 | 4 | 8.5 | 9 |  | 5 |  |  |  |
| 33 | 10.5 | 11.5 | 4.5 | 9.5 | 6.5 | 4 | 8 | 9.5 |  | 5.5 |  |  |  |
| 34 | 11.5 | 10 | 5 | 9 | 7.5 | 6 | 9.5 | 10 |  | 5.5 |  |  |  |
| 35 | 10 | 10.5 | 3 | 9 | 6.5 | 4.5 | 9.5 | 9 |  | 5.5 |  |  |  |
| 36 | 10 | 9 | 5 | 9 | 6.5 | 4.5 |  |  |  |  |  |  |  |
| 37 | 9.5 | 8.5 | 5 | 7 | 7 | 4 |  |  |  |  |  |  |  |
| 38 | 9 | 9 | 5.5 | 6 | 6.5 | 4 |  |  |  |  |  |  |  |
| 39 | 9 | 8.5 | 5 | 6 | 5.5 | 4.5 |  |  |  |  |  |  |  |
| 40 | 7.5 | 8 | 5.5 | 5.5 | 5.5 | 4.5 |  |  |  |  |  |  |  |

**Supplemental Table 1. Clinical EAE scores for all mice used in remission calculations.**
| Day of EAE | Ts EAE 1 | Ts EAE 2 | Ts EAE 3 | Ts EAE 4 | Ts EAE 5 | Ts EAE 6 | Ts EAE 7 | Ts EAE 8 | Ts EAE 9 | Ts EAE 10 | Ts EAE 11 | Ts EAE 12 | Ts EAE 13 | Ts EAE 14 | Ts EAE 15 | Ts EAE 16 | Ts EAE 17 |
| --- | --- | --- | --- | --- | --- | --- | --- | --- | --- | --- | --- | --- | --- | --- | --- | --- | --- |
| 0 | 0 | 0 | 0 | 0 | 0 | 0 | 0 | 0 | 0 | 0 | 0 | 0 | 0 | 0 | 0 | 0 | 0 |
| 2 | 0 | 0 | 0 | 0 | 0 | 0 | 0 | 0 | 0 | 0 | 0 | 0 | 0 | 0 | 0 | 0 | 0 |
| 4 | 0 | 0 | 0 | 0 | 1 | 0 | 0 | 0 | 1 | 0 | 2.5 | 2.5 | 2.5 | 1.5 | 1.5 | 0 | 2 |
| 7 | 0 | 0 | 0 | 0 | 1 | 0 | 1 | 0 | 1 | 0 | 2.5 | 3.5 | 2.5 | 3 | 1.5 | 2 | 4 |
| 8 | 0 | 0 | 0 | 1 | 0 | 0 | 1 | 0 | 1 | 0 | 2.5 | 4 | 4 | 3.5 | 2 | 3.5 | 2.5 |
| 9 | 0 | 1 | 1 | 0 | 1 | 0 | 0 | 0 | 1 | 1.5 | 2.5 | 2.5 | 3 | 4 | 2.5 | 4 | 4 |
| 10 | 0 | 1 | 1 | 0 | 1 | 1 | 0 | 0 | 1 | 1 | 3.5 | 2.5 | 4 | 4 | 4 | 1.5 | 4 |
| 11 | 0 | 1 | 1 | 0 | 1 | 0 | 1 | 1 | 1 | 2.5 | 1 | 2.5 | 1 | 4 | 4 | 2.5 | 4 |
| 12 | 2 | 1 | 1 | 0 | 1 | 0 | 1 | 1 | 1.5 | 2.5 | 2 | 3 | 3.5 | 5 | 2.5 | 3.5 | 4 |
| 13 | 2 | 2 | 2 | 0 | 1 | 0 | 0 | 0 | 1 | 1 | 3.5 | 2 | 1 | 4 | 3.5 | 2.5 | 2 |
| 14 | 2 | 2 | 1 | 1 | 1 | 0 | 1 | 1 | 1.5 | 4 | 2.5 | 1 | 2 | 4 | 4 | 2 | 4 |
| 15 | 2.5 | 2.5 | 2.5 | 0 | 1 | 1.5 | 1 | 1 | 1.5 | 1.5 | 1.5 | 1.5 | 4 | 3.5 | 3.5 | 1.5 | 3 |
| 16 | 2.5 | 4 | 2.5 | 1 | 1.5 | 0 | 1.5 | 0 | 1 | 2.5 | 3.5 | 2 | 4 | 4 | 4 | 4.5 | 3 |
| 17 | 2.5 | 7 | 4 | 0 | 1.5 | 1 | 2.5 | 0 | 2.5 | 2.5 | 3.5 | 4 | 4.5 | 4 | 4.5 | 4.5 | 3.5 |
| 18 | 2.5 | 8 | 4 | 0 | 1.5 | 2.5 | 4 | 2 | 3 | 4.5 | 4 | 4 | 6 | 4.5 | 5 | 5 | 4 |
| 19 | 4 | 7.5 | 7 | 2.5 | 3 | 2.5 | 8 | 1 | 4.5 | 8 | 4 | 4.5 | 4.5 | 5 | 6 | 5 | 4 |
| 20 | 4 | 9 | 7.5 | 0 | 3.5 | 2.5 | 8 | 4 | 4 | 10.5 | 4 | 4.5 | 5 | 5 | 5 | 5.5 | 4.5 |
| 21 | 4 | 8.5 | 10 | 1 | 2.5 | 1 | 8.5 | 5.5 | 1.5 | 13 | 1.5 | 6 | 2.5 | 3.5 | 5 | 6 | 2 |
| 22 | 4.5 | 7 | 10.5 | 1.5 | 2.5 | 1.5 | 8 | 6.5 | 3.5 | 13 | 4 | 8.5 | 3.5 | 3.5 | 6 | 8 | 4.5 |
| 23 | 5.5 | 7 | 9 | 3.5 | 2.5 | 3.5 | 8 | 6 | 3 | 12 | 3.5 | 10 | 5 | 3.5 | 10.5 | 9.5 | 4 |
| 24 | 8 | 7 | 8.5 | 3.5 | 5 | 6 | 6.5 | 6 | 4 | 11 | 4 | 10 | 4 | 2 | 12 | 7 | 5 |
| 25 | 10 | 8 | 8.5 | 6 | 5 | 9 | 5.5 | 6 | 4.5 | 9 | 5 | 9.5 | 6 | 4 | 12 | 7.5 | 5 |
| 26 | 11 | 7 | 7.5 | 7 | 7 | 9 | 6 | 6 | 4 | 10 | 5.5 | 8 | 6.5 | 5 | 12 | 8 | 5.5 |
| 27 | 10 | 6 | 5 | 9 | 7.5 | 10 | 6 | 7 | 7.5 | 9.5 | 6 | 7 | 6.5 | 5.5 | 11 | 4.5 | 5.5 |
| 28 | 9 | 7 | 4 | 7 | 9.5 | 8 | 6 | 7 | 10 | 11 | 5.5 | 5.5 | 8.5 | 6.5 | 9 | 4.5 | 6 |
| 29 | 7 | 6.5 | 6.5 | 9 | 9 | 9 | 7 | 6 | 9 | 8 | 7 | 5.5 | 8.5 | 7.5 | 9 | 5 | 4 |
| 30 | 8 | 7 | 6.5 | 6 | 9 | 7 | 4.5 | 5 | 8.5 | 7 | 7 | 3.5 | 5 | 6.5 | 9 | 4 | 7.5 |
| 31 | 5.5 | 7.5 | 6.5 | 5.5 | 8 | 6 | 4.5 | 5.5 | 9 | 7 | 5 | 5 | 7 | 5.5 | 7 | 4 | 9.5 |
| 32 | 4.5 | 7.5 | 7 | 5.5 | 7.5 | 6 | 4 | 5 | 8 | 6.5 | 5 | 5.5 | 7.5 | 6.5 | 7 | 4 | 9 |
| 33 | 6 | 5 | 6.5 | 5.5 | 7 | 6 | 3.5 | 4.5 | 6 | 7.5 | 5 | 5 | 5 | 4.5 | 6 | 4 | 9 |
| 34 | 4 | 5 | 5.5 | 5 | 8.5 | 5 | 5.5 | 4.5 | 5.5 | 7 | 4.5 | 5 | 4.5 | 3 | 6 | 4.5 | 9 |
| 35 | 3 | 4.5 | 5.5 | 4 | 7.5 | 5.5 | 3.5 | 4 | 5 | 6 | 4 | 5 | 5 | 4 | 7 | 4 | 8.5 |
| 36 | 4.5 | 4.5 | 5.5 | 4 | 7.5 | 5.5 | 3.5 | 4 | 5 |  | 4.5 |  | 4 | 4 |  |  | 9 |
| 37 | 4.5 | 5 | 4 | 4 | 8.5 | 3.5 | 4.5 | 3.5 | 4.5 |  | 4 |  | 4 | 3.5 |  |  | 9 |
| 38 | 4 | 6 | 4.5 | 3.5 | 8 | 4 | 4.5 | 3.5 | 3.5 |  | 4.5 |  | 4 | 4.5 |  |  | 7.5 |
| 39 | 3.5 | 3.5 | 3 | 4 | 9 | 3 | 4 | 4 | 4.5 |  | 4 |  | 4 | 4.5 |  |  | 7.5 |
| 40 | 4.5 | 4 | 3 | 4.5 | 7 | 3 | 5 | 4 | 4 |  | 4 |  | 3.5 | 4.5 |  |  | 7.5 |
| 41 | 4.5 | 5 | 3 | 3 | 9 | 3.5 | 4.5 | 3 | 3 |  | 3 |  | 4.5 | 3.5 |  |  | 6 |

**Supplemental Table 1. Clinical EAE scores for all mice used in remission calculations.** Related to Figure 1E-I. All NI/EAE and *Ts/*EAE mice that had clinical data at least to 10 days post-peak were included. Data are pooled from 4 independent experiments. The “peak” clinical EAE score is highlighted in orange, and the 10 subsequent days are highlighted in yellow.

**Supplemental Table 2. Nanostring nCounter Host Response gene expression summary.** Panel 1) Spleen normalized gene expression of all samples (n=3 NI/EAE, n=3 *Ts/*EAE). Panel 2) Spleen differential expression summary. Panel 3) Inguinal lymph node (iLN) normalized gene expression of all samples (n=3 NI/EAE, n=3 *Ts/*EAE). Panel 4) iLN differential expression summary. Panel 5) Full gene list in Nanostring nCounter Mouse Host Response panel.

**Supplemental Table 3. Microglia RNA sequencing gene expression summary.** Panel 1) Microglia normalized gene expression (n=5 Naïve, n=6 *Ts* 10 DPI, n=4 *Ts* 28 DPI). Panel 2) Differential expression of 10 DPI vs Naïve. Panel 3) Differential expression of 28 DPI vs Naïve. Panel 4) Gene Ontogeny (GO) terms enriched in 10 DPI. Panel 5) GO terms enriched in 28 DPI.

